# Context-Aware Transcript Quantification from Long Read RNA-Seq data with Bambu

**DOI:** 10.1101/2022.11.14.516358

**Authors:** Ying Chen, Andre Sim, Yuk Kei Wan, Keith Yeo, Joseph Jing Xian Lee, Min Hao Ling, Michael I. Love, Jonathan Göke

**Author notes:** contributed equally (Ying Chen, Andre Sim).

## Abstract

Most approaches to transcript quantification rely on fixed reference annotations. However, the transcriptome is dynamic, and depending on the context, such static annotations contain inactive isoforms for some genes while they are incomplete for others.

To address this, we have developed Bambu, a method that performs machine-learning based transcript discovery to enable quantification specific to the context of interest using long-read RNA-Seq data. To identify novel transcripts, Bambu employs a precision-focused threshold referred to as the novel discovery rate (NDR), which replaces arbitrary per-sample thresholds with a single interpretable parameter. Bambu retains the full-length and unique read counts, enabling accurate quantification in presence of inactive isoforms.

Compared to existing methods for transcript discovery, Bambu achieves greater precision without sacrificing sensitivity. We show that context-aware annotations improve abundance estimates for both novel and known transcripts. We apply Bambu to human embryonic stem cells to quantify isoforms from repetitive HERVH-LTR7 retrotransposons, demonstrating the ability to estimate transcript expression specific to the context of interest.

## Introduction

Transcription of DNA is a complex process where multiple alternative RNA transcripts can be expressed from the same gene loci ^1–4^. Transcripts which are derived from alternative gene isoforms can be functionally distinct, making it essential to quantify not just the level of transcription for each gene, but for each individual gene isoform ^5,6^.

The expression levels of these transcripts can be inferred through high throughput sequencing of RNA or cDNA (RNA-Seq) ^7–10^. Sequencing reads are then assigned to known gene isoforms that are catalogued in reference genome annotations. Reference annotations aim to be a comprehensive atlas of an organism’s isoforms that capture all possible tissues and cellular stages. However, due to the dynamic nature of the transcriptome only a fraction of the annotated transcripts are expressed in any given sample, while additional, sample-specific isoforms might be missing from the reference ^11,12^. This particularly impacts transcripts originating from repetitive sequences such as retrotransposons that are challenging to annotate, or cell types such as early embryonic cells that can have a large number of cell-type specific transcripts ^13–15^.

The ability of long read RNA-Seq to generate reads corresponding to full-length transcripts provides an opportunity to discover novel transcripts and thereby enable the quantification of isoform expression using context-specific annotations ^16^. Tools such as FLAIR ^17^, TALON ^18^, or StringTie2 ^19^ have been developed for transcript discovery from long read RNA-Seq and have been shown to identify novel transcripts even in well annotated genomes. However, RNA degradation, sequencing, and alignment artefacts can introduce false positive transcript candidates and impact quantification ^20^.

To deal with possible false positive novel transcripts, existing methods rely on user defined thresholds such as the minimum read count to filter novel transcript candidates ^17–19^. However as these parameters are dependent on sequencing depth, the same threshold can generate vastly different results across multiple samples ^21–23^. This can partially be addressed with additional thresholds such as a minimum relative isoform expression or transcripts-per-million. While these thresholds correct for sequencing depth, they are still influenced by aspects such as expression level or number of isoforms per gene and may erroneously filter out valid novel transcript candidates. Therefore, as none of these thresholds provide an intuitive way of controlling false positive transcript candidates, bounding the error rate of the resulting novel transcript set remains a challenge.

To overcome these limitations we have developed Bambu, a method for multi-sample transcript discovery and quantification. Bambu estimates the likelihood that a novel transcript is valid, allowing the filtering of transcript candidates with a single, interpretable parameter, the novel discovery rate (NDR), that is calibrated to guarantee a reproducible maximum false discovery rate across different samples and analyses. Bambu then employs a statistical model to assign reads to transcripts that distinguishes full-length and non full-length (partial) reads, as well as unique and non-unique reads, thereby providing additional evidence from long read RNA-Seq to inform downstream analysis. We demonstrate that the NDR implemented in Bambu enables transcript discovery with greater accuracy and with a wider dynamic range compared to existing tools. Additionally we show that Bambu’s context-specific annotations improve quantification even for annotated transcripts, and that the ability to track unique and full-length reads reduces the impact of inactive transcripts. We apply Bambu to long read RNA-Seq from human embryonic stem cells (hESCs) and illustrate the ability to quantify individual isoforms from highly repetitive HERVH transposons with full-length read support. Together, Bambu addresses the limitation of static reference annotations while providing a quantitative measure of confidence for novel transcripts, enabling the comprehensive, context-aware quantification of individual isoforms from long read RNA-Seq.

## Results

### Bambu: context aware quantification of long read RNA-Seq data

Bambu consists of 4 steps: Firstly, a probabilistic model is employed to correct junction alignments using reference annotations, genome sequence, and features obtained from the data (see Methods). Corrected reads that use the same splice junctions are summarised into *read classes* (Figure 1a). In the second step, read classes from all samples are combined and a cross-sample NDR is calculated. Read classes below the specified NDR threshold are treated as novel transcripts, resulting in context-specific reference annotations (Figure 1b). Thirdly, each read class is assigned to compatible transcripts allowing for inexact matches to account for possible alignment errors (Figure 1c). In the fourth step, transcript expression estimates are obtained with an expectation-maximisation (EM)^24^ algorithm for each sample using the same set of context-specific transcript annotations. Bambu estimates expression levels using reads that are uniquely assigned to a single transcript (*unique* reads) as well as reads which are assigned to multiple transcripts. Bambu then provides final expression estimates that include the number of *full-length* reads and *unique* reads that support each transcript. While Bambu models expression jointly using all reads, the individual contributions from each group are tracked to provide an intuitive measure of evidence for each transcript and gene to support their expression in the samples of interest (Figure 1d). Bambu is available through Bioconductor, it runs with a single command and a single, interpretable transcript discovery parameter, and can efficiently be applied to a large number of samples (Supplementary text section 9, Supplementary Text Figure 4, Supplementary Text Table 4-5), thereby greatly simplifying the quantification with context-specific annotations from long reads.

**Figure 1.**
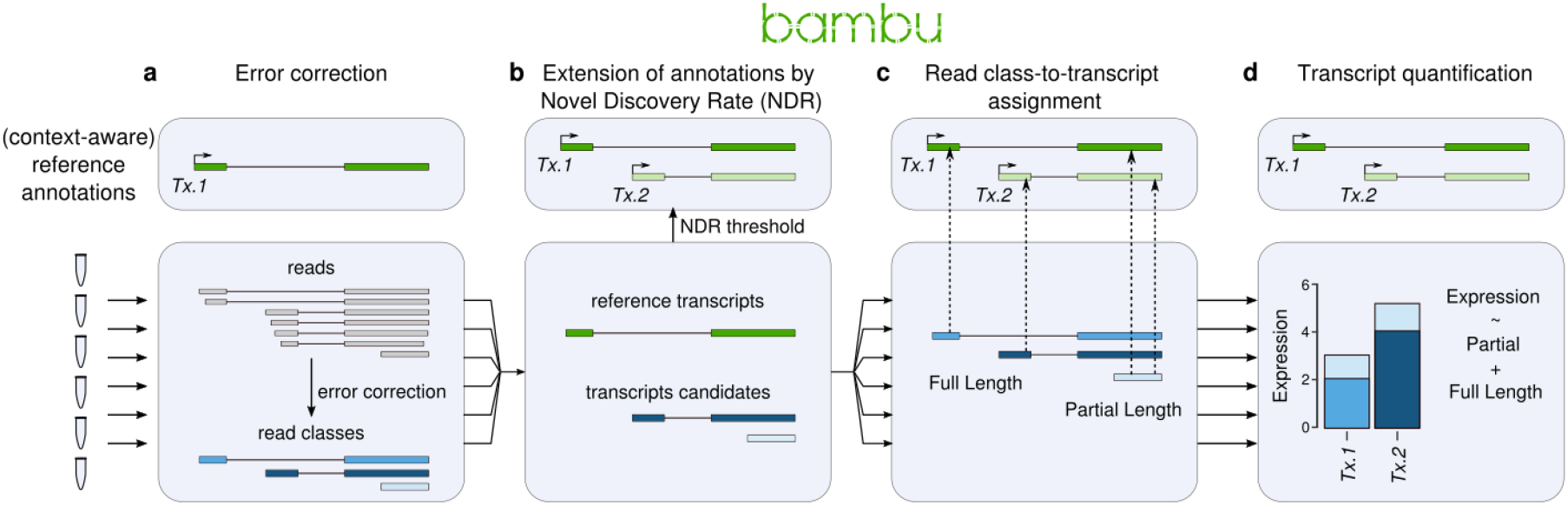
Bambu enables simultaneous transcript discovery and quantification from Nanopore RNA-Seq data. Schematic illustration on how Bambu performs transcript discovery and quantification on Nanopore RNA-Seq data in four steps **(a)** For each sample, Bambu performs error correction on splice junctions of the aligned reads using input annotations **(b)** Performs transcript discovery jointly across samples at a given novel discovery rate (NDR) threshold and extends the input annotations with the retained novel transcripts **(c)** Assigns the read classes to transcripts in the extended annotation and categorises them as having full-length or partial overlaps **(d)** Performs probabilistic transcript quantification based on the read class to transcript assignment

### Bambu infers a transcript probability score to accurately rank novel transcript candidates

In order to obtain a single parameter for transcript discovery, Bambu extracts nine different features that summarise read count, read alignment, and read sequence characteristics and trains a supervised machine learning model that infers the probability for each read class to be a valid transcript (transcript probability score) (Figure 2a, see Methods and Supplementary Text Section 1). The multi-sample transcript score is then defined as the probability that a transcript is valid in at least one sample (see Methods and Supplementary Text Section 3), enabling the ranking of transcript candidates across multiple samples without the need to apply any threshold on individual samples.

**Figure 2.**
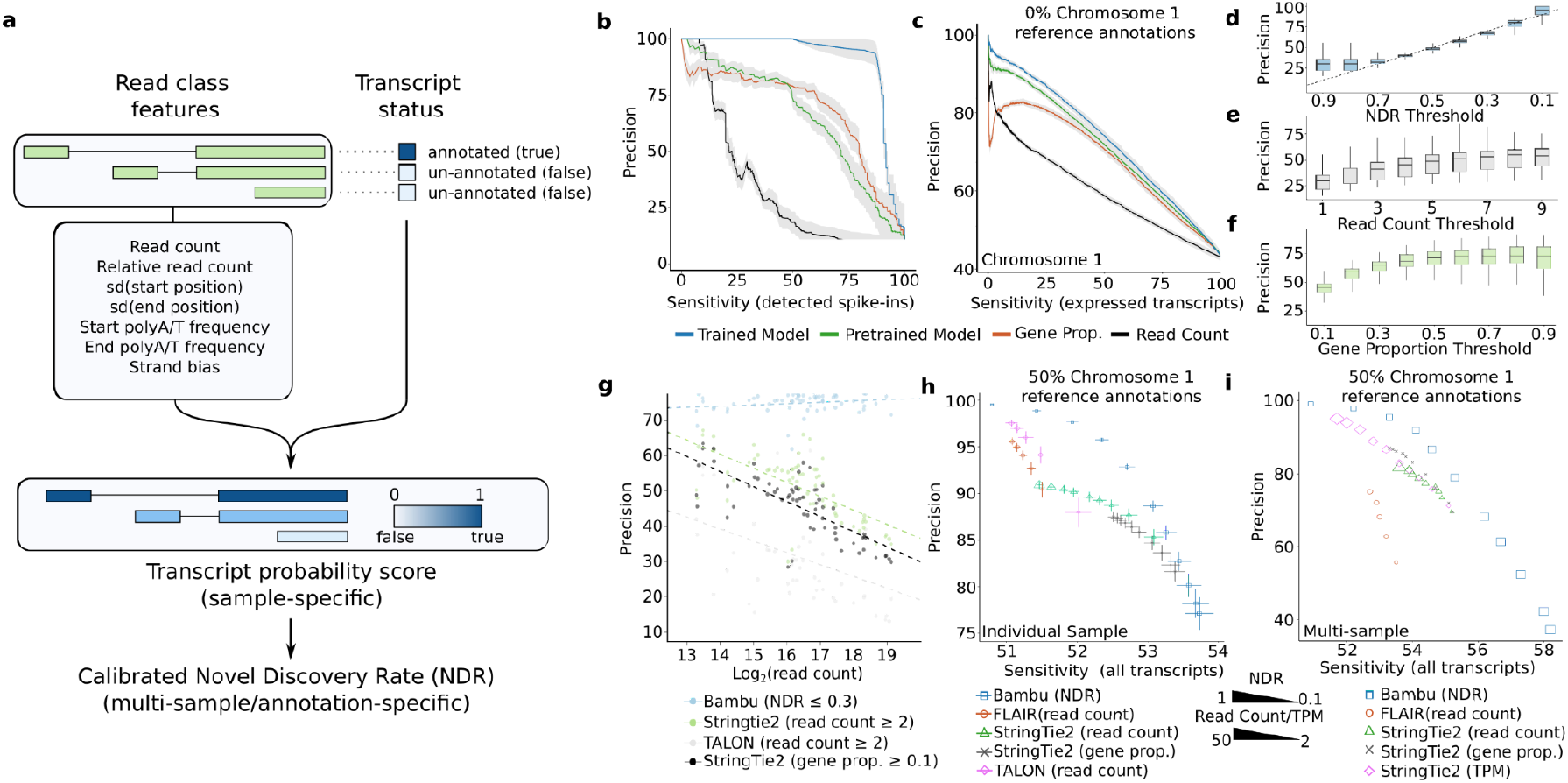
calibrated machine learning full-length transcript classifier improves transcript discovery accuracy. **(a)** The schematic of transcript discovery steps performed by Bambu where 1) a machine learning model is trained on nine different features from read classes features to predict if a read class represents a full-length transcript, 2) the transcript probability score predicted in the first step is re-calibrated to a novel discovery rate across multiple samples **(b)** Precision Recall curves for the performance of transcript discovery for spike-in data. Each line shows the average precision and recall of the model on all SG-NEx spike-in data after being trained on a subset of the spike-in data (blue), predictions from the generic model (green), or when read count (black) or gene proporton (red) is used alone as a classifier. The grey shaded area represents the mean +/- SE of the precision for each line **(c)** Precision Recall curves for the performance of transcript discovery for human chromosome 1 data. All read classes aligned to chromosome 1 were used to test the model. The model was trained using read classes from all other chromosomes (blue), the generic model (green), or when read count (black) or gene proporton (red) is used alone as a classifier. The grey shaded area represents the mean +/- SD of the precision for each line. Sensitivity is measured as the percentage of true transcripts out of all expressed transcripts. **(d-f)** Boxplot showing the median, upper and lower quartiles, and 1.5 x interquartile range of the precision of chromosome 1 read classes passing varying (d) NDR, (e) Read Count and (f) Gene Proportion thresholds when removing chromosome 1 from the reference annotations across all SG-NEx samples. Precision is measured as the percentage of read classes that completely match splicing junctions for human chromosome 1 transcripts of all read classes above the threshold **(g)** The resulting precision for each annotation output when a fixed threshold is used to reclassify the annotations found on human chromosome 1. Each dot colour triplicate represents the same SG-NEx sample processed by either: Bambu with a NDR threshold of 0.3 (blue), StringTie2 with a read coverage threshold of 2 (green), TALON with a read count threshold of 2 (grey) and StringTie2 with a gene proportion threshold of 0.1 (black). Dotted lines represent a fitted linear regression for each tool **(h)** The average sensitivity and precision of transcript discovery on core SG-NEx samples with 50% of human chromosome 1 annotations randomly removed. Each tool is displayed at several different parameter thresholds: Bambu (blue) with NDR thresholds varying between 1 and 0.1, FLAIR (red), Stringtie 2 (green) and TALON (pink) with read count/coverage thresholds varying between 2 and 10, with 4 additional thresholds for StringTie2 at 15, 20, 30 and 50. StringTie2 was also thresholded using gene proportion (black). Horizontal error bars represent the mean +/- SD of the sensitivity and vertical error bars represent the mean +/- SD of the precision. Sensitivity is measured as the percentage of reported transcripts out of all annotated transcripts from chromosome 1, a minimum of 50% sensitivity is expected as 50% of the annotations are provided. **(i)** The measured sensitivity and precision of transcript discovery when combining HepG2 samples, with 50% of human chromosome 1 annotations randomly removed. Each tool is displayed at several different parameter thresholds: Bambu (blue) with NDR thresholds varying between 1 and 0.1, FLAIR (red) and Stringtie 2 (green) with read count/coverage thresholds varied between 2 and 10, as well as 4 additional thresholds for StringTie2 at 15, 20, 30 and 50. Stringtie 2 was also run with a read count threshold of 2 and a varying TPM threshold between 2 and 50 (pink). StringTie2 was also thresholded using gene proportion (black). Horizontal error bars represent the mean +/- SD of the sensitivity and vertical error bars represent the mean +/- SD of the precision.

Bambu uses existing annotations as training data, and learns a model for each sample assuming that the majority of valid transcripts are annotated. However, for genomes which are poorly annotated or samples with a very high number of expected novel transcripts, a pretrained model can be used. Bambu includes a model that was trained on Nanopore RNA-Seq data from a human cell line (see Methods). However, for cases where the pretrained model is not suitable due to species and technology differences, Bambu provides the option to train a new model on related data with comprehensive annotations that is more appropriate for the sample of interest (see Supplementary Text Section 5 for additional information).

To evaluate the performance of the transcript score to identify valid transcripts, we first applied Bambu on sequin spike-in RNAs ^25^ where the ground truth is known, using Nanopore long read RNA-Seq samples from the Singapore Nanopore Expression (SG-NEx) project^26^. On these data, the transcript score shows a high level of precision, outperforming the baseline parameters of read counts and relative isoform gene expression (gene proportion) as parameters for transcript discovery (Figure 2b). While the sample-specific model shows the highest performance, the pre-trained model still outperforms baseline statistics (Figure 2b). Next we tested the performance for transcript discovery on the human chromosome 1 for which annotations were removed before running Bambu. The results similarly show that both the sample-specific and the pre-trained model have a higher level of precision compared to the baseline parameters, and it is consistent when applied on single samples or multiple samples (Figure 2c, Supplementary Figure 1a). We repeated this analysis using Arabidopsis and PacBio RNA-Seq data, which further confirmed that the sample-specific model and the pre-trained model show better performance than read count or gene proportion (Supplementary Figure 1b-c). The TPS was even able to rank candidates that were lowly expressed or which consisted only of a single isoform, something not possible when using baseline read count and gene proportion thresholds (Supplementary Figure 1d-e).

To test the robustness of the supervised approach used in Bambu when annotations are incomplete, we trained a model providing 25%, 50%, 75% and 100% of human annotations respectively. Even with 50% of annotations we observe that the model trained in Bambu shows a high level of accuracy (Supplementary Figure 1f, Supplementary Text Table 1). When less than 50% of annotations are known, the model still generates accurate predictions, however a pretrained model using more complete annotations is able to provide improved performance (Supplementary Figure 1f). To catch such a scenario in practice, Bambu automatically infers the completeness of reference annotations and recommends using a pre-trained model when the expected fraction of missing transcripts is higher than 50% (Supplementary Figure 1g-h, see Methods and Supplementary Text Section 5). Even when trained on very poorly annotated genomes (25% of annotations), the sample-specific model shows a higher overall accuracy than read count (Supplementary Figure 1f, Supplementary Text Table 1). Together, these results indicate that the transcript score predictions in Bambu provide a robust and accurate way to rank and identify valid transcripts on both well annotated and poorly annotated genomes (Figure 2c, Supplementary Figure 1f, Supplementary Text Table 1).

### The Novel Discovery Rate (NDR) replaces arbitrary per-sample thresholds with a single, interpretable, and comparable parameter

In order to make the transcript discovery parameter interpretable and comparable across samples, we define the novel discovery Rate (NDR) as the fraction of unannotated transcripts among all transcripts with an equal or higher transcript score. The NDR can be interpreted as an upper limit on the false discovery rate (or 1-precision) under the assumption that reference annotations are complete: a NDR of 0.1 indicates that at least 90% of transcripts with a similar score or higher are annotated, thereby providing an intuitive estimate of precision. As the expected number of novel transcripts differs depending on the completeness of the annotations, the same NDR can correspond to different levels of precision across different species or when alternative annotations are used. However, in practice most analyses are done within the same species and annotations where the number of expected novel transcripts is similar, in which case the same NDR threshold guarantees a similar FDR and precision across independent analysis.

To evaluate this, we identified novel transcripts with different NDR thresholds for all samples in this study. Here we provide the human genome annotations without chromosome 1 during transcript discovery, using annotations from chromosome 1 as ground truth to estimate the precision of transcript discovery. A comparison of the observed precision for different NDR thresholds confirms that it is indeed well calibrated (Figure 2d). We find that the same NDR threshold provides a very similar level of precision across all samples, whereas an equivalent read count or gene proportion threshold results in a wide range of precision (Figure 2d-f). Furthermore, unlike thresholds such as read count, TPM, or relative read count that are used by other methods, the NDR provides a continuous metric that is linearly related to the expected precision (Figure 2d-f). This property enables transcript discovery across the complete dynamic range of precision, facilitating either conservative but accurate extension of annotations in the case of well annotated genomes, or more sensitive transcript discovery for genome annotation of species or samples with many unknown transcripts. To optimise results for each analysis where samples have varying levels of annotation completeness, Bambu infers an analysis-specific default NDR threshold using the estimate of the fraction of missing transcripts (see Methods). Using the pretrained model, Bambu is able to estimate the fraction of missing annotations accurately in ONT, PacBio and other species (Mouse and Arabidopsis) data (Supplementary Figure 1g, Supplementary Text Table 2). This dynamic default threshold is calibrated to ensure high levels of precision, however it can be changed for more sensitive transcript discovery. As the NDR is calculated on the multi-sample transcript probability score, it replaces sample-specific thresholds with a single, interpretable, and comparable parameter for transcript discovery.

### Bambu achieves a higher dynamic range and accuracy compared to existing methods

Next we benchmarked Bambu against FLAIR, TALON and StringTie2 using the SG-NEx long read RNA-Seq samples. First we compared the impact of transcript discovery parameters on precision, confirming that read-count based parameters generate vastly different levels of precision for identical thresholds (Figure 2g). In contrast, the same NDR threshold provides a comparable precision across independent samples (Figure 2g). Next, we evaluated the precision and sensitivity to identify valid multi-exon transcripts when 50% of the transcript annotations were removed from chromosome 1 in the reference (see Methods). A comparison of the different methods demonstrates that Bambu achieves higher precision at a comparable sensitivity when samples are analysed individually (Figure 2h). The same results are obtained when applied to spike-in transcripts and when no annotations were provided (Supplementary Figure 1i-l). Furthermore, Bambu enables transcript discovery with a wider dynamic range of precision compared to all other tools (Figure 2h).

A unique feature of Bambu is its ability to analyse results across multiple samples with a single, calibrated threshold. When all SG-NEx samples are jointly analysed, Bambu maintains the ability to perform transcript discovery across the full dynamic range of precision, while other methods that use non-calibrated thresholds applied to each sample showed a smaller range of precision or sensitivity (Figure 2i). This property is particularly relevant for well annotated genomes (e.g. human), where high sample numbers with high sequencing depth and using read-count-based thresholds would otherwise result in high numbers of novel transcripts that might impact downstream quantification. Together these results show that Bambu is more accurate, provides a wider range of precision and is the only method where the precision can directly be controlled with a single transcript discovery parameter.

### Context-specific annotations improve transcript quantification

After transcript discovery, Bambu estimates transcript expression using the full length and partial length read classes for all samples. We first compared the quantification in Bambu without transcript discovery (NDR=0) to existing quantification-only methods Salmon^10^, NanoCount^27^, featureCounts^28^, and LIQA^29^ using the sequin spike-in RNAs with complete reference annotations. We find that quantification with Bambu on spike-in RNAs had comparable or better performance to the existing quantification-only tools (Figure 3a-b).

**Figure 3.**
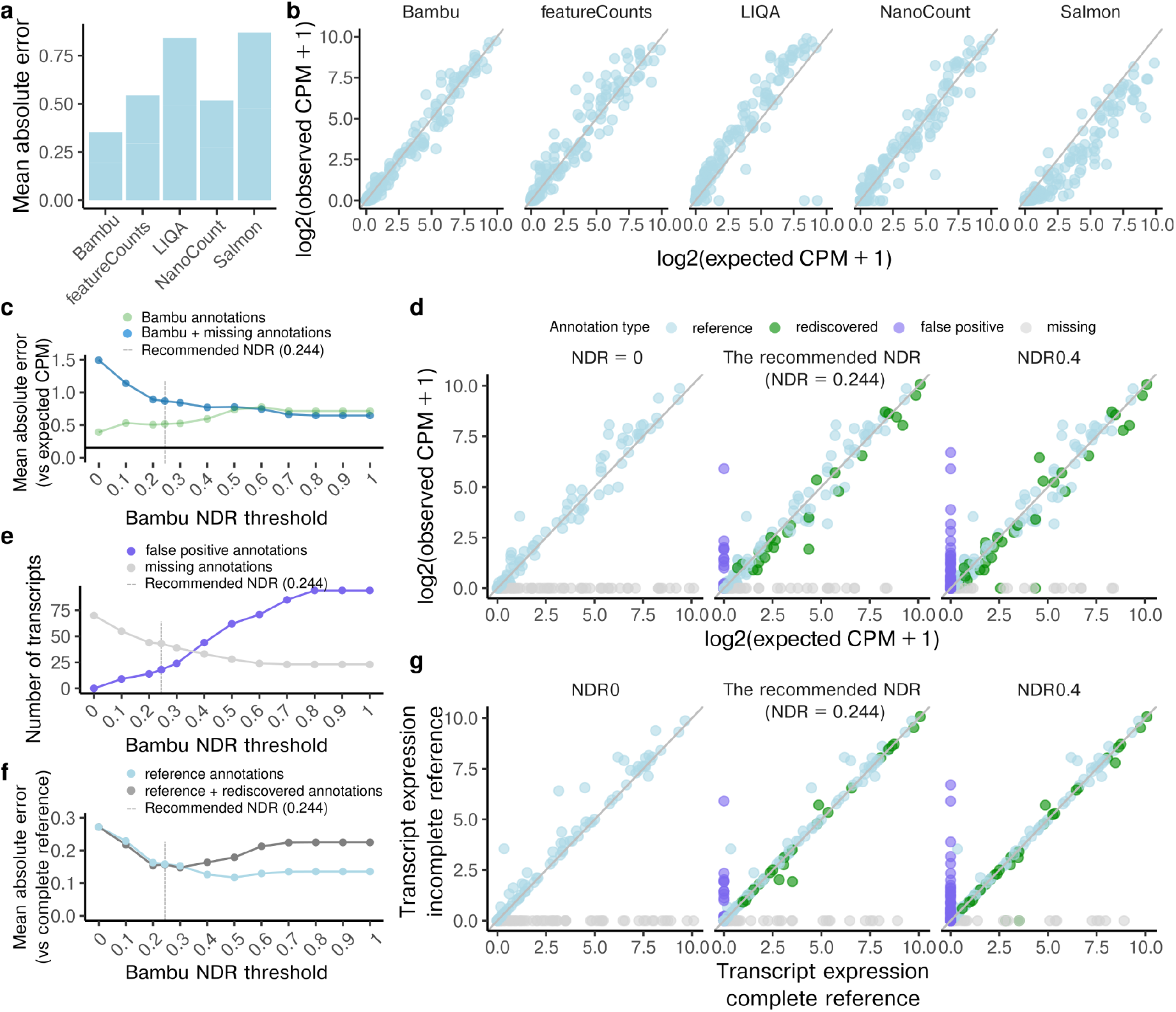
Transcript quantification on spike-in data shows improvement with varying novel discovery rates. **(a)** The mean absolute error between the log2 normalised transcript abundance estimates and log2 normalised expected spike-in abundance when applying Bambu, featureCounts, LIQA, NanoCount, and Salmon with full sequin annotations. **(b)** Scatterplots between log2 normalised transcript abundance estimates and log2 normalised expected spike-in abundance when applying Bambu, featureCounts, LIQA, NanoCount, and Salmon with full annotations. **(c)** The mean absolute error between the log2 normalised transcript abundance estimates and log2 normalised expected spike-in estimates using partial sequin annotations when applying Bambu with varying novel discovery rate thresholds (NDR), for Bambu annotated transcripts, including annotations that are present in the reference (partial) sequin annotations, the annotations that have been artificially removed and rediscovered by Bambu, and also the false positive annotations discovered by Bambu (green), plus the annotations that are artificially removed from the partial annotation and remained missing after transcript discovery, i.e., missing annotations (blue). The grey dotted line indicates the recommended NDR by Bambu (0.244) **(d)** A log2 normalised scatterplot of Bambu estimates when partial annotations are used against true concentration levels for transcripts that are present in the partial annotations(light blue), transcripts that have been artificially removed from the reference and rediscovered by Bambu (green), false positive transcripts (purple), transcripts that have been artificially removed from the reference and remained missing after Bambu discovery (grey) when applying Bambu: without transcript discovery (NDR = 0),with default recommended NDR (0.244), and with a more sensitive NDR (0.4) **(e)** The number of missing (grey) and false positive (purple) transcripts using partial sequin annotations when applying Bambu with varying NDR thresholds. The grey dotted line indicates the recommended NDR by Bambu (0.244) **(f)** The mean absolute error between the log2 normalised spike-in transcripts estimates with partial and full annotation when applying Bambu with varying novel discovery rate thresholds (NDR), for transcripts that are present in reference (light blue), transcripts that are present in reference and those rediscovered annotations (dark grey). The grey dotted line indicates the recommended NDR by Bambu (0.244) **(g)** A scatterplot of log2 normalised Bambu estimates when partial annotations are used against when full annotation is used for transcripts that are present in the reference (light blue), transcripts that have been artificially removed from the reference and rediscovered by Bambu (green), false positive transcripts (purple), transcripts that have been artificially removed from the reference and remained missing after Bambu discovery (grey) when applying Bambu: without transcript discovery (NDR = 0),with default recommended NDR (0.244), and with a more sensitive NDR (0.4)

Next, to evaluate the impact of transcript discovery on quantification, we specified different NDR thresholds and compared the accuracy of abundance estimates for spike-in RNAs with partially missing annotations. Quantification after transcript discovery in Bambu (NDR>0) allows the abundance estimation for missing gene isoforms, reducing the overall estimation error (Figure 3c-d, Supplementary Figure 2a-c, Supplementary Figure 3a). More sensitive transcript discovery will increase the number of false positive transcripts, highlighting the importance of choosing a threshold that is appropriate for the analysis (Figure 3e).

Our results further suggest that transcript discovery also stabilises abundance estimates for isoforms which are already present in the reference annotation, leading to more consistent results on the same data and higher reproducibility with increasing NDR (Figure 3f-g, Supplementary Figure 2d-e, Supplementary Figure 3b). The same observation is made when the extended annotations from Bambu are used with other quantification-only tools, where we observe that transcript discovery reduces the quantification error of annotated transcripts (Supplementary Figure 4–5).

A comparison of quantification after transcript discovery in Bambu with other transcript discovery methods shows higher variation across the tools (Supplementary Figure 2). These results reflect the differences quantification methods, but also in the extended annotations that differ for each tool and which are defined by tool-specific default parameters, precision, and sensitivity, further highlighting the impact of transcript discovery on quantification and emphasising the value of the single parameter in Bambu that enables the control of false positive transcripts and their impact on quantification.

### Full-length and unique read support provide additional evidence that transcripts are expressed in the samples of interest

Even in long read RNA-Seq data, reads that match multiple transcripts are still frequently present (*non-unique* reads: 13.8 - 49.5%, Supplementary Figure 6a). Similar to existing methods, Bambu uses an EM algorithm to probabilistically assign non-unique reads to transcripts. While this approach has been previously demonstrated as effective ^27^, there is no guarantee that transcripts which are only supported by non-unique reads are indeed expressed in the samples of interest. To address this, we calculate for each read if it matches a complete transcript (full-length), and if it can be uniquely assigned to a single transcript (unique) (see Methods). Bambu then provides estimates of the full-length and unique read support in addition to the total abundance estimation in counts per million (CPM) for each gene isoform. When we compared transcript expression estimates across biological replicates, we found that transcripts with a higher number of unique or full-length reads show higher correlation, suggesting that they provide additional information that is not captured by the total CPM (Figure 4a, Supplementary Figure 6b-e).

**Figure 4.**
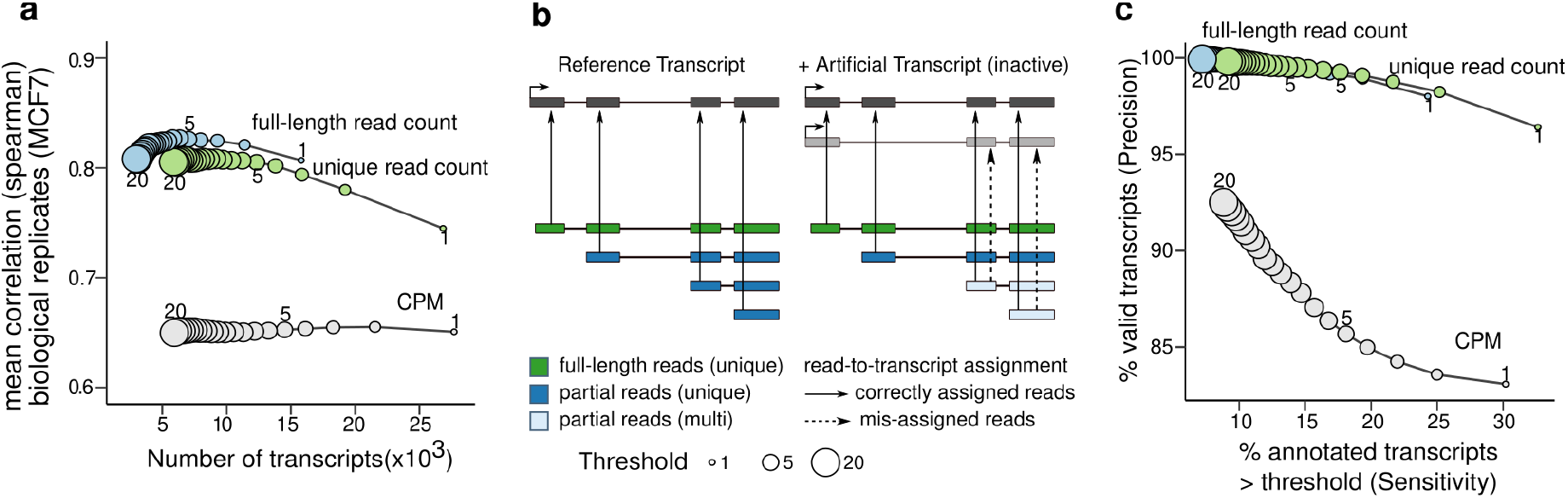
Full-length and unique read support provide evidence on expressed transcripts. **(a)** The average spearman correlation between the transcript abundance estimates for MCF7 replicates generated using direct cDNA and the number of expressed transcripts when different filtering methods and thresholds were applied to transcripts. Filtering is based on mean CPM or unique read count support being greater than the given threshold across the replicates **(b)** Full-length, unique read and partial reads and the potential read-to-transcript assignment for each of these read types **(c)** The sensitivity and precision of using full-length, unique read and CPM thresholds to filter out false positive transcripts overlapping with highly abundant isoforms that have no unique or full-length reads support at varying filtering thresholds from 1 to 20 on Hct 116 samples. Filtering was based on the average values across replicates being not lower than the threshold.

To evaluate if CPM, unique, and full-length read counts can be used as evidence that a transcript is expressed in the samples of interest, we included artificial isoforms containing a unique splice junction (exon skipping) in the reference annotation prior to quantification (Figure 4b). These artificial isoforms are not expressed, however, due to probabilistic read-assignment and approximate read matching in Bambu, they can still be estimated to be active (Figure 4c, Supplementary Figure 6f). Increasing the CPM threshold reduces the number of such transcripts, indicating that read-misassignment mostly affects transcripts with lower overall read count, and suggesting that a basic expression filter already provides some evidence that a transcript is expressed (Figure 4c). Using the presence of unique reads or full-length reads further improved the ability to identify expressed transcripts, achieving a higher precision at the same level of sensitivity when compared to the basic CPM expression filter (Figure 4c, Supplementary Figure 6g). While Bambu considers all reads to obtain the CPM estimate, the ability to track full-length and unique reads enables Bambu to quantify transcript abundance in the presence of unexpressed reference annotations, as well as providing an additional layer of evidence to inform downstream analysis or follow up experiments.

### Bambu enables the quantification of retrotransposon-derived transcripts

Among the most difficult genes to quantify are those that are derived from retrotransposons, as they are highly repetitive and often not accurately annotated. One of the most striking examples of retrotransposon expression is the HERVH-LTR7 (Human endogenous retrovirus subfamily H-Long Terminal Repeat) family which has been reported to be a highly specific marker of pluripotency in hESCs ^30–33^. The human genome contains several thousand annotated HERVH-LTR7 elements. However, the number of expressed elements is expected to be much smaller ^34^. To test if Bambu can enable the detailed reconstruction of HERVH-derived transcripts purely from RNA-Seq data, we analysed the hESC samples from the SG-NEx data. We observed a significant enrichment of repetitive sequence in novel genes (p < 0.001), with HERVH-LTR7 being the dominating repeat family (Figure 5a). In total, 242 genes (encompassing 464 transcripts) are transcribed from HERVH, 64 of them contributing to 90% of the HERVH RNA in hESCs, suggesting that only a small minority of HERVH elements are actually transcribed (Figure 5b, Supplementary Figure 7, and Supplementary Table 1). These HERVH-derived genes are supported by full-length reads, and they generate distinct transcripts with alternative splicing patterns that are unique to each locus, suggesting that they might be functionally distinct (Figure 5c-f). Quantifying transcript expression in hESCs without the extended annotations from Bambu results in an overestimation of existing transcripts such as *ESRG*, where the majority of reads originate from previously undescribed isoforms (Figure 5d, Supplementary Figure 7c). Together, these results illustrate how context-aware quantification with Bambu reduces quantification error while enabling the estimation of individual retrotransposon-derived isoform expression from long read RNA-Seq without any additional experimental or computational requirements.

**Figure 5.**
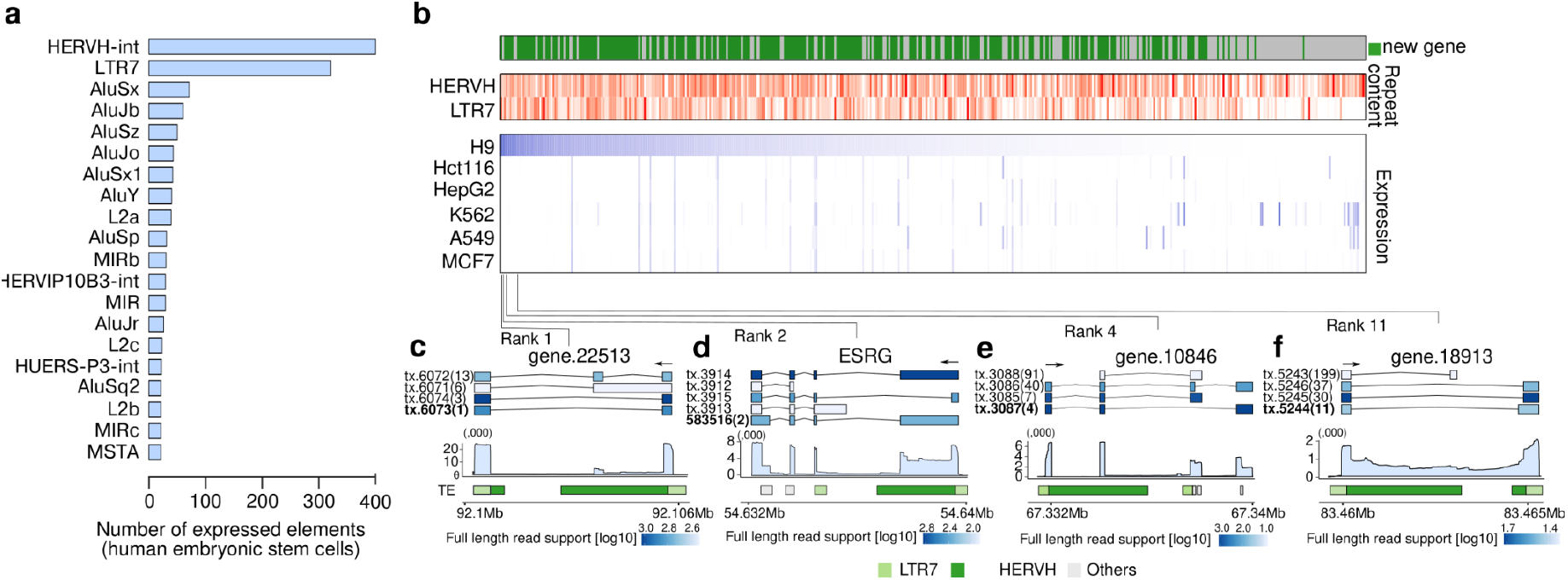
Bambu enables the discovery and quantification of highly repetitive genes. **(a)** Repeat families ranked by the number of expressed elements identified in the human embryonic stem cell cell line (H9) **(b)** Overview of the top 100 expressed novel and annotated transcripts that overlap with the HERVH-LTR7 retrotransposon in hESC cell line. For novel transcripts, we only include those with high overlap in novel exons. Top: Fraction of overlap with HERVH and LTR7. Bottom: Expression estimate in H9 hESCs and in the 5 SG-NEx cancer cell lines **(c-f)** Illustrations of highly expressed transcripts in H9 hESCs that originate from HERVH-LTR7 repeats that show distinct splicing patterns and transcript sequences. Top: transcript annotation colored by estimated full-length read support, with ranks in expression highlighted inside the bracket. Middle: mean read coverage for the specified genomic ranges for each selected gene. Bottom: show repeat masker colored by HERVH (green), LTR7 (light green), and all other repeat types (light grey).

## Discussion

While static reference annotations simplify the quantification of transcript expression, they do not necessarily reflect the samples or analysis of interest, with unknown transcripts continuing to be discovered even in well annotated species ^21–23^. To address this, we developed Bambu, a method that enables quantification of transcript expression with annotations that are inferred specifically for the context of interest using long read RNA-Seq. The accuracy of transcript discovery is influenced by the parameter thresholds that are applied to identify novel transcripts ^35,36^. Bambu employs a machine learning approach to control the false positive rate using a single transcript discovery parameter (the NDR). In contrast to parameters such as read count, relative expression and TPM, the NDR ranks transcript candidates by their probability of representing a valid full-length transcript. Therefore, unlike arbitrary thresholds for parameters used in other tools ^17–19^, a more stringent NDR threshold guarantees higher precision while providing an upper bound on the FDR. This is especially relevant for well annotated genomes or analyses which involve high numbers of samples where precision is more important than sensitivity to obtain accurate annotations and quantification results. However, even analyses where a high number of novel transcripts are expected will benefit from this property as the NDR identifies the most precise set of annotations even when more sensitive thresholds are selected.

In most cases, the expected number of novel transcripts is unknown before transcript discovery. To avoid arbitrary and often inappropriate default thresholds, Bambu estimates the fraction of annotations that are missing in the sample, which is then used to recommend a suitable NDR for the analysis. Therefore, unlike other tools that rely on fixed default thresholds ^17–19^, Bambu infers a threshold to achieve high precision in transcript discovery and accurate quantification.

While Bambu uses annotations for training the transcript prediction model, splice site correction and NDR calibration, it also works well on poorly annotated genomes and those without any annotation. In these scenarios a pretrained model predicts the transcript probability score without the need for reference annotations. Bambu provides a model trained on nanopore RNA-Seq data from human cell lines, however, for samples with vastly different genomes or alternative sequencing technologies, Bambu includes the option to pre-train a model on a similar dataset. Without reference annotations the NDR is not estimated, and transcripts are instead ranked by the uncalibrated transcript probability score from the pretrained model. While the transcript score is not calibrated to be comparable across analysis, its ability to rank transcript by precision is identical to the NDR, making it possible to discover novel transcripts with low false discovery rates even in the absence of annotations.

Very sensitive transcript discovery provides more novel transcript candidates, however, we observe that the increasing complexity of annotations can impact quantification due to reads which are assigned to multiple transcripts. Long-read RNA-Seq is able to generate reads which match full-length transcripts, thereby providing evidence that transcripts are present in the samples of interest ^21,35–37^. In Bambu we use all reads for quantification, while providing the full-length read count estimate for each transcript. However, the classification of reads as full-length in Bambu is influenced by both sequencing and alignment errors as well as RNA degradation, with non-unique full-length reads still being ambiguous. Approaches have been developed to enrich full-length reads experimentally ^38^, or selectively analyse full-length reads during quantification ^39^. These approaches are more effective in identifying which transcripts are truly expressed, however, as they significantly reduce the number of reads used for quantification, the overall abundance estimates are likely to be less accurate. Here we find that non-unique full-length reads only represent a minor fraction of reads, suggesting that for most genes, Bambu’s approach provides accurate quantification while still preserving the key information about full-length reads for each transcript.

The ability to perform context-specific quantification is particularly relevant for applications where novel transcripts are expected. When applying Bambu to hESCs, we discovered many retrotransposon-derived genes that are missing from current annotations, with full-length reads providing an additional layer of evidence. Furthermore, Bambu enables applications in other areas, including identifying and quantifying individual fusion transcripts when reads are aligned to the breakpoint-corrected genome, or combined with tools for detection of RNA modifications from direct RNA-Seq data, Bambu can provide insights into epitranscriptomic changes at novel transcripts. Yet, Bambu is not just limited to long read RNA-Seq. With a small set of representative long read samples, Bambu can be used to generate context-specific annotations to further improve quantification from large-scale short read data sets. In conclusion, Bambu simplifies transcript discovery and quantification across multiple samples to a single command with a single parameter, while improving accuracy and interpretability of expression estimates, suggesting that quantification with context-specific annotations could become a routine approach to analysing transcript expression with long reads.

## Methods

### Transcript Discovery and Quantification with Bambu

Bambu performs both transcript discovery and quantification with the following steps:

#### (1) Construction of Read classes

##### 1.1 Alignment error correction

Firstly, to utilise long reads which can have high noise during basecalling resulting in errors in splice-junction alignments, we trained a machine learning model with scalable tree boosting to effectively predict the probability of a true splicing junction. This was performed for every 5’ and 3’ splice site considering information from splicing junctions within 15 base pair (bp) distance. The model is trained using R XGBoost function with default parameters ^40^ and uses 5’ and 3’ distances between read alignments and annotation, strand information, the presence of splice motifs, and read support relative to the reference splicing junctions as features. Splicing junctions that are predicted to be true will be noted as high confidence junctions and maintained, while junctions which are predicted to originate from mis-alignments are noted as low confidence junctions and will be corrected to the closest reference splice site within 10 bp, or kept with the original alignment if no reference splice site is within this range.

##### 1.2 Construction of Read Classes

Here we assume that reads with identical splice patterns originate from the same transcripts. Under this assumption, we summarise all reads spanning the same (error-corrected) junctions into *read classes* (*RC*). The start coordinate of the read class is defined as the position that includes 80% of all read start positions for this read class, the end coordinate is similarly defined to include 80% of the read end positions for each read class. Only read classes with high confidence junctions (and optionally read classes which are unspliced) are retained for transcript discovery, whereas all read classes are used for quantification. Novel read classes are matched to genes on the basis of exon overlap with reference annotations. Read classes which do not overlap with reference annotations are classified as belonging to a novel gene, assigned a new gene id, and grouped together with other read classes which they overlap.

#### (2) Feature extraction and Model Training

To predict the probability that a read class represents a valid transcript, Bambu uses a supervised machine learning approach using XGBoost. Firstly, Bambu extracts 9 features from the read classes: read counts-per-million (referred to as read count), the proportion of contribution to the total read count of its corresponding gene (referred to as gene proportion), the proportion of reads mapping to the strand with higher read count, the standard deviation of supporting reads’ 5’ and 3’ ends positions, and the frequency of A/Ts in the first and last 20 bp of the read class as features. (See Supplemental Test for additional details). Bambu then represents each read class *i* from sample *j* with a feature vector *x_i,j_* and an associated binary class label *y_i,j_*. that indicates if a read class is a valid transcript:

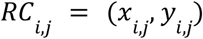

with *x_ij_* = {*read count, relative read count*,…}
and *y_i,j_* = {1 *if RC_i,j_ is a valid transcript*; 0 *otherwise*}.

The probability that a read class represents a valid transcript (the Transcript Probability Score) for read class *RC_i,j_* is then defined as

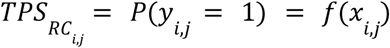

Here, *f*(*x*) represents the sample-specific model that is trained for each sample *j* using scalable tree boosting with the XGBoost (with default parameters and 50 stumps used in the decision trees). During training, *y_i,j_* is defined using the set of all reference transcript annotations *T*:

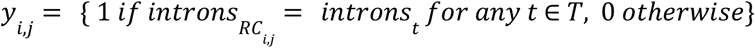

To reduce the noise during training, we exclude any read class with only single read count and read classes that do not overlap with any reference annotation (novel genes). Because Bambu relies on splice-junction coordinates to define *y_i,j_* during training, single exon read classes are excluded. However, Bambu optionally trains a separate model for read classes which consist of only a single exon, and which can predict the TPS for these transcript candidates (option min.txScore.singleExon = 0 in Bambu argument opt.discovery, see Supplementary Text Section 4).

##### 2.1 The pre-trained model in Bambu

Training of *f*(*x*) requires annotations to define class labels. When less than 1000 annotated transcripts are expressed in the sample (very poorly annotated genomes), training a sample-specific model is not supported due to insufficient training data from annotated transcripts. In this case, a pre-trained generic model can be used. Model pre-training can be done with a well annotated genome that is closely related to the genome of interest, or a pre-trained model from Bambu can be used. The default pre-trained model is based on the SGNex_HepG2_directRNA_replicate5_run1 sample from SG-Nex resource which is applicable to any species with similar characteristics as the human genome (Supplementary Text Figure 1.b). The pre-trained model is also used to recommend a transcript discovery threshold, see Methods (below), Supplementary Text and the Bambu online documentation for additional details.

###### 2.1.1 Automatic recommendation procedure to use a pre-trained model

More complete annotations will result in a more accurate transcript discovery model learned by Bambu. As the pre-trained model is assumed to be learned on very well annotated genomes, it shows higher accuracy compared to a sample-specific model when annotations are poor or the data is very noisy. If the estimated fraction of missing transcripts is below 50% (see below for details on the estimation of the fraction of missing transcripts), Bambu will recommend to use a pre-trained model, resulting in a mean improved performance for these samples even when the minimum annotation threshold to train a sample-specific model is satisfied (Supplementary Figure 1h).

##### 2.2 Multiple Sample Transcript Discovery

In Bambu, transcript discovery is performed across all samples *j* = 1,…, *N*. Here we define the Transcript Probability Score for a read class *i* as the probability that it is a valid transcript in at least 1 sample:

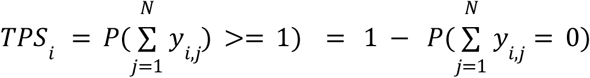

To avoid that technical replicates inflate the multi-sample TPS, we provide a greatest lower bound using the maximum TPS for read class *i* across all samples:

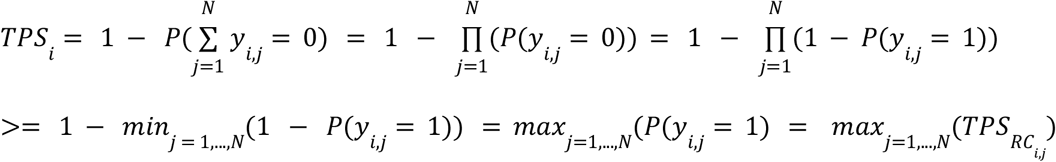

For a comparison of alternative definitions of the multi-sample TPS, please see the Supplementary Text 3. In the case of a single sample, *TPS_RC_i__* = *TPS_RC_i,j__*, therefore we use TPS to refer to the multi-sample TPS for simplicity.

#### (3) Prediction of novel transcripts using the Novel Discovery Rate (NDR)

##### 3.1 Transcript discovery threshold using the TPS

To identify novel transcripts, the TPS for each read class is estimated. Novel transcripts are then identified based on a TPS threshold *p*:

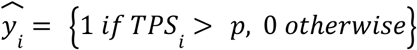

For each threshold *p*, the number of read classes predicted to be valid transcripts is defined as

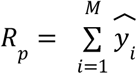

##### 3.2 Limitations of sample-specific thresholds

Please refer to Table 1 for definition of error rates in the following section. The TPS can be used to rank transcripts by their probability of being valid. The rank of read class *i* is defined as the number of read classes with less or equal TPS:

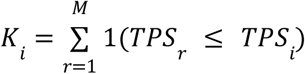

with the function r providing the map between *i* and *K_i_*:

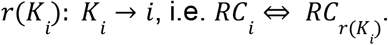

**Table 1:** Definition of error rates during transcript discovery

|  | Invalid transcript | Valid transcript |  |
| --- | --- | --- | --- |
| Predicted valid transcript | $V^{FP} = V - V^{TP}$ | $S + V^{TP}$ | $R$ |
| Predicted invalid transcript | $U + U'$ | $T + U'$ | $N - R$ |
| | $n_0 - n'$ | $n_1 + n'$ | $M$ |
| **observed:**<br>$V$: number of non annotated read classes with $TPS_i > p$<br>$S$: number of annotated read classes with $TPS_i > p$<br>$U$: number of non annotated read classes with $TPS_i < p$<br>$T$: number of annotated read classes with $TPS_i < p$<br>$R$: number of read classes with $TPS_i > p$<br>$M$: Total number of read classes<br>$n_0$: total number of non-annotated read classes<br>$n_1$: total number of annotated read classes<br>**Not observed:**<br>$V'$: number of non annotated valid read classes with $TPS_i > p$<br>$U'$: number of non-annotated valid read classes with $TPS_i < p$<br>$n'$: total number of non-annotated valid read classes |  |  |  |
| **Table 1:** Definition of error rates during transcript discovery |  |  |  |

The number of true positive (valid) novel transcripts associated with threshold *p* is then defined as

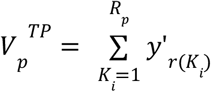

with *y′_i_* indicating the unknown true class label of read class *i*. The number of false positive novel transcripts associated with threshold *p* is defined as

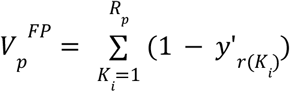

The False Discovery Rate (FDR) associated with *p* is then defined as

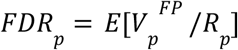

Furthermore, we define the Valid Discovery Rate (VDR) as the expected number of valid novel transcripts associated with *p*:

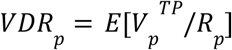

The VDR can be interpreted as the expected fraction of missing transcripts in the annotations. Since the scale of the TPS can differ with each new analysis based on the samples and annotations that are provided during model training (i.e. *f_j_*(*x*) ≠ *f*(*x*)’ for any two different analysis samples *j* and *j′*), the same threshold *p* can lead to results with a different false discovery rates (*FDR_p_j__* ≠ *FDR_p_j__*) making the selecting of a meaningful and consistent threshold *p* difficult in practice (the same limitation applies to all sample-specific parameters such as read count). To address this in Bambu, users can specify the expected novel discovery rate. Bambu then selects the optimal threshold p for the specified NDR.

##### 3.3 The Novel Discovery Rate: definition and calculation

To obtain a calibrated score that provides comparable and interpretable thresholds, we define the Novel Discovery Rate (NDR) as the expected number of non-annotated transcripts:

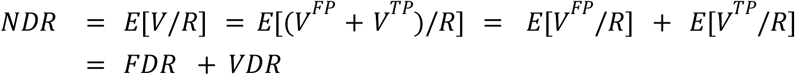

Therefore, the NDR can be interpreted as an upper limit of the FDR for very well annotated genomes (FDR>VDR) or as an upper limit of the VDR for poorly annotated genomes (*FDR* ≪ *VDR*) (see below for additional information on interpretation of the NDR). To select the threshold p that maximises the number of novel transcripts for the specified target NDR, we first calculate the number of novel transcripts associated with each candidate threshold p’ as:

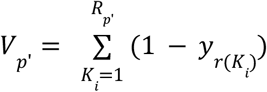

With *y_i_* indicating if read class *RC_i_* matches any annotation that is provided (see above). The observed *NDR^o^_p’_* for p’ is then calculated as:

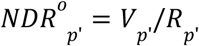

We then select the threshold p which corresponds to the largest number of novel transcripts such that the observed NDR is below or equal the target NDR:

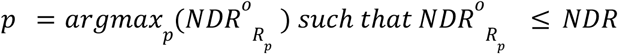

The maximum number of novel transcript candidates for the target NDR is then

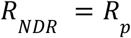

Each read class is then associated with an optimal threshold *p* ≤ *TPS_i_* such that *NDR_P_* ≤ *NDR_i_*, and the NDR associated with each read class is then estimated as

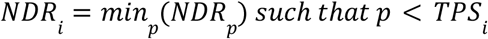

Therefore, *NDR_i_* ≤ *NDR_i_, if TPS_i_ ≥ TPS_i’_*

##### 3.4 Properties and Interpretation of the NDR

The interpretation of the NDR differs for very well annotated genomes (transcript discovery) and poorly annotated genomes (genome annotation):

###### 3.4.1 Transcript discovery for well annotated genomes (FDR>VDR)

For well annotated genomes such as the human genome, it is expected that most transcripts are annotated (FDR>VDR). In this scenario, the NDR provides an upper limit of the FDR:

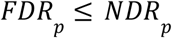

Here, the NDR enables the selection of a threshold p that maximises the number of novel transcripts for a desired maximum FDR. Unlike sample-specific threshold, the NDR ensures that the maximum FDR is comparable across samples or analyses:

*FDR_p_j__* ≤ *NDR* ≥ *FDR_p_j__* with *p_j_* and *p_j_′* corresponding to the optimal threshold for the target NDR in sample *j* and *j′* respectively. Since VDR>0 in most cases, the NDR is a conservative estimate of the FDR, with the true *FDR* ≪ *NDR*.

###### 3.4.2 Genome annotation (transcript discovery for poorly annotated genomes)

For poorly annotated species (for example from newly assembled genomes), it is expected that a large fraction of novel transcripts are missing, in which case FDR≪NDR. In this scenario the NDR does not provide a meaningful upper limit of the FDR. However, the FDR is expected to increase dramatically once the NDR exceeds the expected VDR. To avoid this when Bambu is used for genome annotation, the NDR can be used as an a upper limit on the expected fraction of novel transcripts:

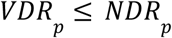

In this scenario, the NDR enables users to select a threshold for transcript discovery that ensures that the maximum number of transcripts is selected such that the expected VDR for high quality annotations is not exceeded, thereby preventing over-annotation with false positive transcripts. Here the NDR is also a conservative estimate of the VDR to ensure that the number of false positive transcripts is minimised.

##### 3.5 Automatic estimation of fraction of missing transcripts

Bambu automatically estimates the fraction of missing annotations (1-completeness of annotations) at a maximum FDR of 0.1 to (1) recommend a default NDR threshold and (2) suggest the use of a pre-trained model for increased accuracy (see above for additional details). To estimate the completeness of the annotations that are provided, the TPS from the pre-trained model is used. During model training, it is assumed that a complete set of annotations is provided, which ensures that 1 - NDR provides an approximation of the precision. Based on this assumption, Bambu identifies the TPS threshold 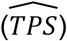 from the pre-trained model corresponding to a precision of 0.9 (i.e., NDR = 0.1, across the SG-NEx data, the mean TPS at this NDR threshold is 0.891 with a SD of 0.065 suggesting the relationship between the TPS and NDR is robust). Bambu then infers the completeness of annotations as the fraction of novel transcripts among all transcripts with 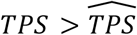 when the pre-trained model is applied. While the Bambu default pre-trained model is based on human RNA-Seq data, any pre-trained model can be used to estimate the completeness of annotations.

##### 3.6 Estimation of a dynamic default NDR

The completeness of the reference annotations should be considered when choosing a NDR threshold: For well annotated genomes, a stringent NDR is recommended, whereas in the case of largely unannotated genomes, a larger fraction of novel transcripts is expected, and a more sensitive NDR threshold is recommended. To moderate the impact of the number of reference annotations on transcript discovery when the NDR is not explicitly specified, Bambu uses a dynamic default NDR which corresponds to a expected maximum False Discovery Rate of 10% for the samples and annotations of interest. To achieve this, Bambu uses the estimated fraction of missing annotations at a FDR of 10% as the default NDR. We would like to note that this is a conservative estimate to minimise the number of false positives. For use cases when sensitivity is most important and higher false discovery rates are acceptable (e.g. genome annotation), this will be an underestimate, and we recommend manually specifying the desired NDR.

#### (4) Additional filtering and final annotation output of transcript discovery

##### 4.1. Additional filtering

Bambu provides additional user-defined filters for which read classes will be considered as novel transcripts. In particular, read classes that (1) are possible degradation or sequencing artefacts can be removed (remove.subsetTx, default: TRUE), (2) have a read count below a minimum threshold low (min.readCount, default = 2), (3) have a gene read proportion below a minimum threshold (min.readFractionByGene, default = 0.05). Furthermore, the minimum number of samples which have to pass the min.readCount and min.readFractionByGene filters can be adjusted. Bambu also allows the specification of the minimum distance to annotated transcripts for read classes (min.exonDistance) to be considered as novel, and the minimum overlap (min.exonOverlap) to merge novel unspliced transcript candidates with annotations. Read classes that pass all the filters and which have a NDR below the (user defined or default dynamic) NDR threshold will then be retained as novel transcript candidates (novel read class candidates). All read classes that pass and that do not pass these filters will be retained and used for quantification.

##### 4.2. Integration with reference annotations

Bambu compares all novel read class candidates with the reference annotations. Read classes which have similar exon junctions compared to reference annotations will not be considered novel transcripts (see online documentation for additional distance thresholds that can be specified). Remaining novel read class candidates will then be included as new transcripts in the reference annotations that are provided (if any are provided). Novel transcripts which overlap with any existing gene are assigned to this gene id, and novel transcripts which overlap with any other novel transcript that is assigned to an annotated gene are iteratively assigned to the same gene id. Bambu then classifies novel transcripts of annotated genes according to the overlap with reference annotations as containing new first exons, new last exons, or new internal exons. Overlapping novel transcripts which are not assigned to any annotated gene will be grouped as novel genes and assigned a novel gene id. Bambu returns the combined reference annotations and novel transcript (referred to as the extended annotation).

#### (5) Quantification

##### 5.1 Read class to transcript assignment

Using the extended reference annotations, Bambu then assigns each error-corrected read class to a set of reference transcripts based on the compatibility of splice junctions and the distance to the most similar reference transcripts. To avoid mis-assignments due to incorrect start and end sites, the first and last exons are optionally excluded. Quantification is performed separately for each sample using the same set of extended annotations. In the following we refer to transcripts as *t_i_*, with *i* = 1,.., *N*, using *j* as the index for reads, read classes or equivalent read classes (see below for definition).

###### 5.1.1 Compatibility of read classes with transcripts and genes

Read classes are assigned to all transcripts and genes which are *compatible:*

###### 5.1.1.1 Transcript compatibility

For a read class *RC_j_* to be considered compatible with transcript *t_i_*, Bambu requires that all splice junctions of *RC_j_* are similar to a continuous set of splice junctions from *t_i_*. By default, Bambu allows for up to 35 bp distance for each exon such that a read class is considered compatible with a transcript. Read classes which are not compatible with any transcript are considered *incompatible* and not considered for transcript quantification. All incompatible read classes are still assignable to genes, which allows us to use them for gene quantification.

###### 5.1.1.2 Gene compatibility

For a read class *RC_j_* to be considered compatible with gene *g_j_*, Bambu requires that any exon of *RC_j_* overlaps by at least 35 bp with any exon from gene *g_i_*. Read classes which are incompatible with all transcripts, but compatible with genes are still used to obtain more accurate gene expression estimates (see below for additional details).

###### 5.1.2 Definition of full length (equal) and partial read classes

Read classes for which (a) all splice junctions are present in a transcript and (b) which are compatible with this transcript are considered to be *equal* to that transcript. All reads that correspond to equal read classes are counted as full-length reads. Compatible read classes which are not equal to any transcript are considered to represent non-full length (*partial*) reads.

###### 5.1.3 Definition of equivalence read classes (equiRCs)

**Table 2:**
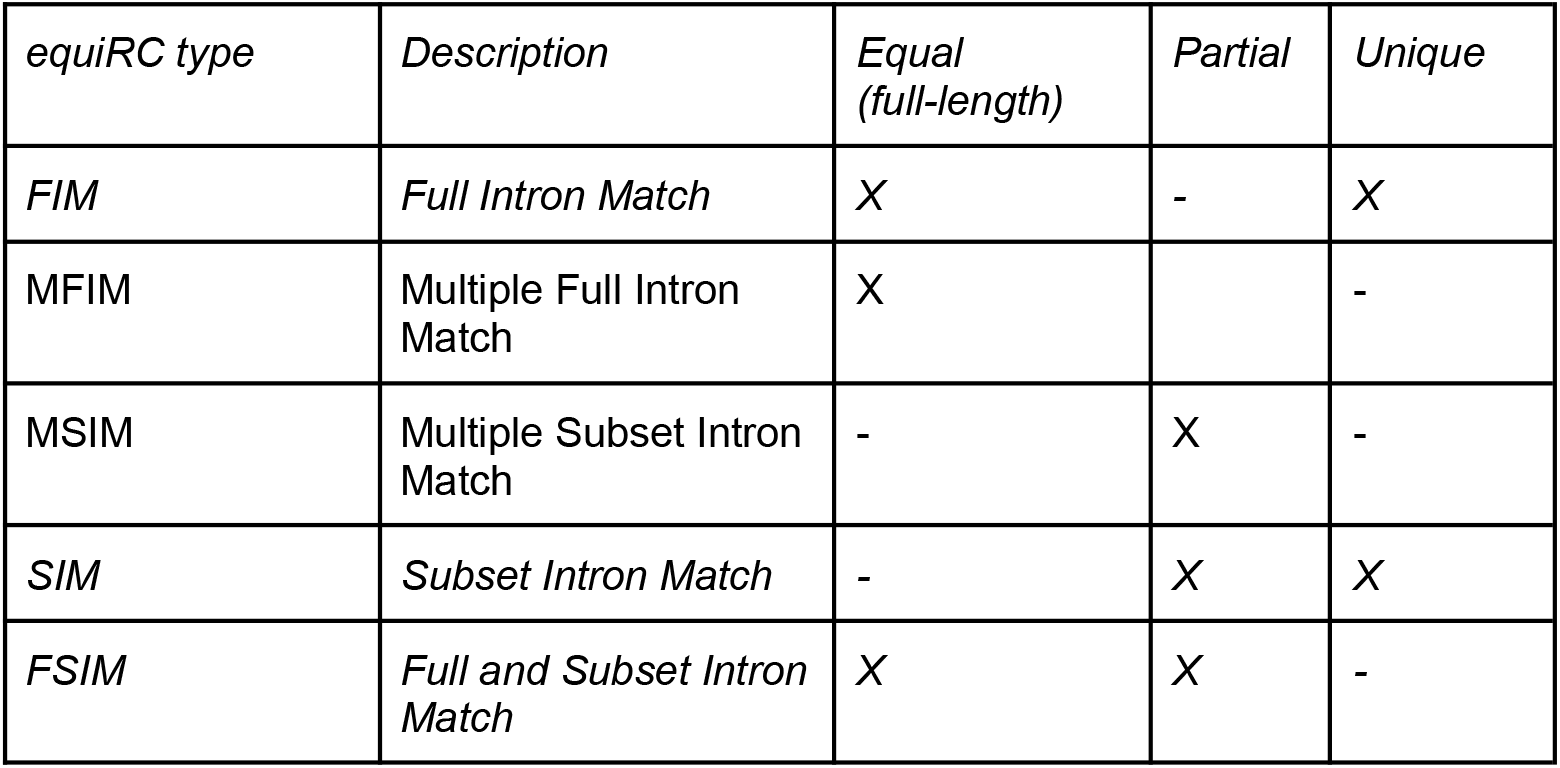
Definition of equivalence read class types

| *equiRC type* | *Description* | *Equal*<br>*(full-length)* | *Partial* | *Unique* |
| --- | --- | --- | --- | --- |
| *FIM* | *Full Intron Match* | X | - | X |
| MFIM | Multiple Full Intron Match | X |  | - |
| MSIM | Multiple Subset Intron Match | - | X | - |
| *SIM* | *Subset Intron Match* | - | X | X |
| *FSIM* | *Full and Subset Intron Match* | X | X | - |

Read classes which are compatible with the same set of transcripts will be summarised into *equivalent read classes* (equiRCs). To leverage the advantage of long read RNA-Seq to generate full-length reads, equal and partial read classes are summarised as different equiRCs. For each equiRC *j*, we observe the number of reads that form this read class (*n_j_*), and the set of transcripts *I_j_* that are compatible.

With this definition, equiRCs can be summarised into five categories (Table 1, Supplementary Figure 5): 1) Full Intron Match equiRC (FIM): equiRCs equally aligned to a unique transcript; 2) Subset Intron Match equiRC (SIM): equiRCs partially aligned to a unique transcript; 3) Multiple Full Intron Match equiRC (MFIM): equiRCS equally aligned to multiple transcripts (due to the existence of very similar transcripts where only first or last exons are different); 4) Full and Subset Intron Match equiRC (FSIM): equiRCS equally aligned to a transcript while partially aligned to one or more longer transcripts; this occurs when transcripts are subsets of other (longer) transcripts, and; 5) Multiple Subset Intron Match equiRC (MSIM): equiRCs of fragmented reads that aligned partially to multiple transcripts.

##### 5.2 Generative model

We followed the conventional generative model used for transcript quantification with specific changes to allow quantification of total expression and full length and unique reads from long read RNA-Seq data ^7,9,10^:

**Table 3.** Description of mathematical notations for transcript quantification. The realisations of these variables are denoted with a prime symbol (′) on top.

| Mathematical notations | Descriptions |
| --- | --- |
| $r$ | read index |
| $i$ | transcript index |
| $j$ | equivalence read classes (equiRC) index |
| $M$ | total number of transcripts |
| $K$ | total number of equivalence read classes (equiRC) |
| $N$ | total number of reads, i.e., sequencing depth |
| $\tau_i(\tau_i^f, \tau_i^p, \tau_i^u)$ | true read count for isoform $i$, superscripts $f, p, u$, represent true |
|  | full-length, partial length, and unique alignment read count respectively |
| $R_{ri}^t$ | the event that whether a read $r$ originates from transcript $i$: takes a value of 1 when it happens and 0 otherwise (the realisation value) |
| $R_{rj}^{rc}$ | the event that whether a read $r$ matches a equiRC $j$ |
| $R_{rij}^{rc,t}$ | the event that whether a read that is assigned to equiRC $j$ originates from transcript $i$ |
| $S_i^t$ | the set of reads that originate from transcript $i$ |
| $S_j^{rc}$ | the set of reads that are assigned to equiRC $j$ |
| $S_{ij}^{rc,t}$ | the set of reads in equiRC $j$ being generated from isoform $i$ |
| $T_j^{rc}$ | the set of transcripts that are compatible with equiRC $j$ |
| $RC_i^t$ | the set of equiRCs which are compatible with transcript $i$ |
| $a_{ij}$ | the conditional probability that a read $r$ matches to a equiRC $j$ conditioning on the read $r$ originates from transcript $i$ |
| $\theta_i$ | global true relative expression of isoform $i$ |
| $n_{ij}$ | unobserved number of reads in equiRC $j$ originate from isoform $i$ |
| $n_j$ | number of reads in in equiRC $j$ |
| $I_{ij}^f (I_{ij}^u)$ | an indicator variable takes a value of 1 when reads in read class $j$ that originate from transcript $i$ are full length reads (unique reads) and 0 otherwise |

Bambu estimates the read count *τ_i_* for each isoform *i* = 1,…, *M* in the sample. We assume that each read *r* = 1,…,*N* originate from a single transcript *i*, which we describe as 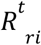:

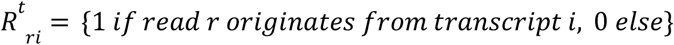

with 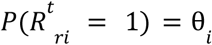. describing the probability that a read *r* originates from transcript *i*, which we also refer to as the relative transcript abundance, and *S^t^_i_*. describing the set of reads that originate from transcript *i*. Then *τ_i_* is defined as:

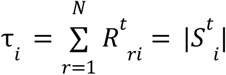

with the expected value for *τ_i_* obtained as:

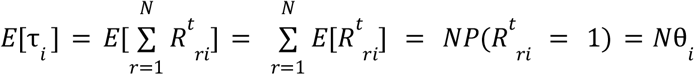

Bambu summarises reads that have the same set of compatible transcripts into equivalent read classes (equiRCs). The assignment of a read r to equiRC *j* is then described as:

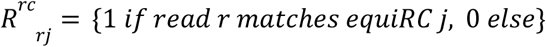

with *j* = 1... *K*. Here we refer to the set of reads that are assigned to equiRC *j* as *S^rC^_j_* with

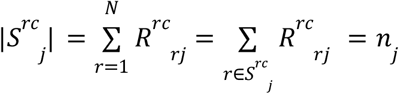

The set of transcripts that are compatible with equiRC *j* is described as *T^rc^_j_* and the set of equiRCs which are compatible with transcript *i* are described by *RC^t^_i_*. The observed value of *R^rc^_rj_*, is always conditional on the (unknown) true read to transcript assignment *R^t^_rj_*. Here we define the conditional probability that a read *r* matches read class *j* given that it originates from transcript *i* as *a_ij_*.

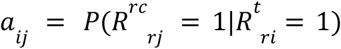

with 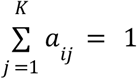 for each transcript *i*. Without prior information, *a_ij_* is assumed to be equally distributed among the set of equiRCs that are compatible with transcript *i* (*RC^t^_i_* with |*RC^t^_i_*| = *K_i_*), and otherwise: 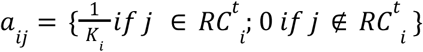.

For each read *r*, 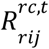 denotes if a read that is assigned to equiRC *j* originates from transcript *i*:

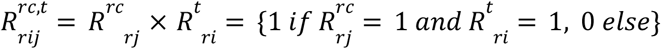

We can then calculate the probability as

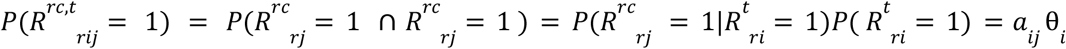

given 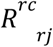 is dependent on 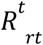.

The set of reads in equiRC *j* being generated from isoform *i* is then denoted as *S^rc,t^_ij_*, with the number of reads in equiRC *j* that originate from isoform *i* defined as *n_ij_*:

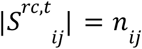

As *n_ij_* is a sum of i.i.d. Bernoulli random variables 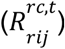, *n_ij_* follows a binomial distribution:

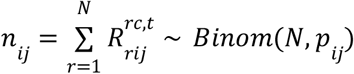

with 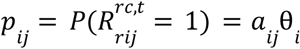.

As Bambu works at the level of read classes for increased computational efficiency, first *n_ij_* is inferred (see below) and the read count for each isoform *i, τ_i_* is then calculated as

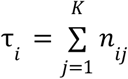

##### 5.3 Parameter estimation using Expectation Maximisation (EM)

To estimate the transcript abundance, we use an Expectation Maximisation (EM) algorithm, that iteratively optimises the likelihood of the relative transcript abundance parameter (θ = {θ_1_,θ_2_,… θ_*M*_}) given the observed read-to-read class assignments 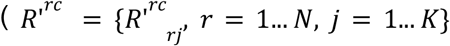, with 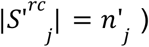 and the latent data that describes for each read *r* the transcript that generated this read 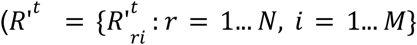, with 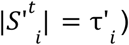. In practice Bambu works at the level of read classes, therefore the latent data used in Bambu’s EM is the read assignment within each read class 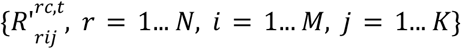, with 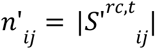 and 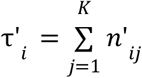.

The complete likelihood function can be written as:

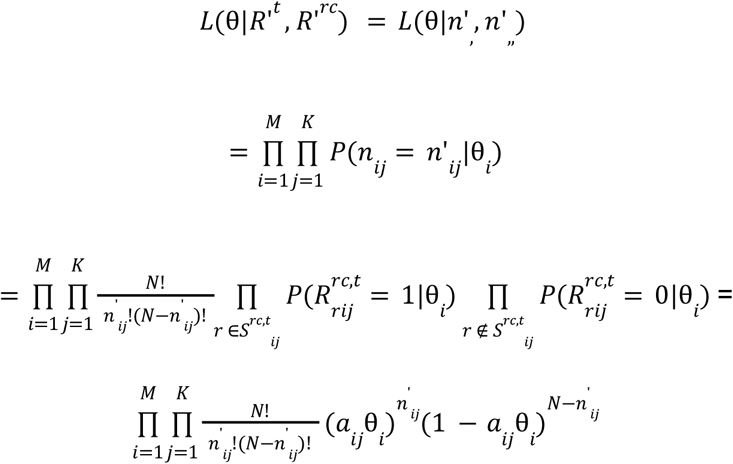

###### Expectation Step

In the *k^th^* expectation step (E-step), the latent data 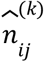 is estimated as the conditional expected value of *n_ij_* given the observed data *R′^rc^* and 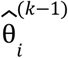:

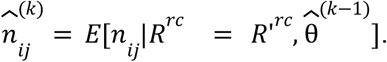

Since we observe the read to read class assignment (*R′^rc^*), we estimate the conditional expected value for *n_ij_* as

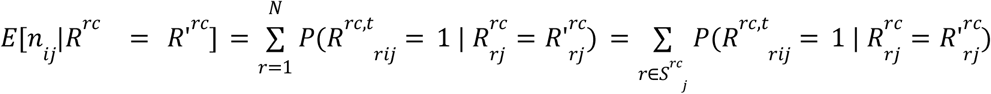

Using the following 2 relations (1) 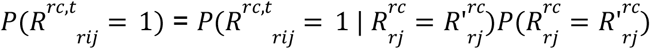 and (2) 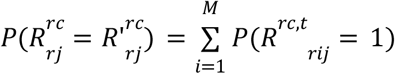 we obtain:

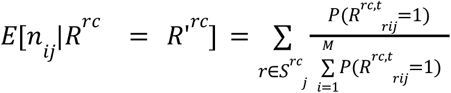

Therefore

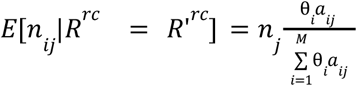

During the E-Step, we use the estimated values 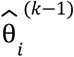 from the previous ((*k* – 1)) iteration (with 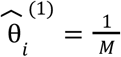), the observed read count for equiRC *j*(*n′_j_*), and the predefined values for *a_ij_* to calculate the estimated value 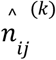 for the kth iteration:

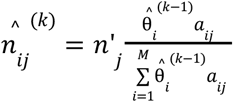

With the estimated transcript abundance for iteration *k* being calculated as 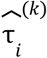:

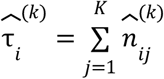

###### Maximisation Step

In the Maximisation Step (M-Step), we obtain the maximum likelihood estimate for the unknown parameter θ given the estimate for the latent data 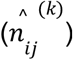:

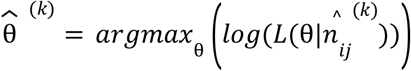

The maximum likelihood estimate can be obtained as follows:

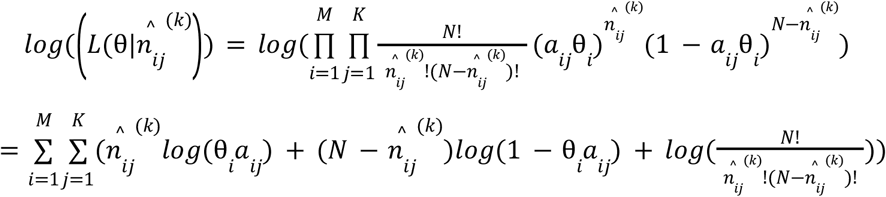

Taking derivative with respect to *θ_i_*:

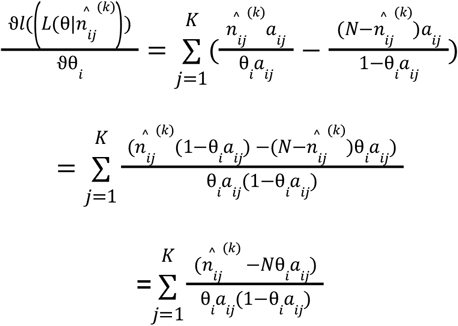

By setting the above equation to 0, therefore, we have

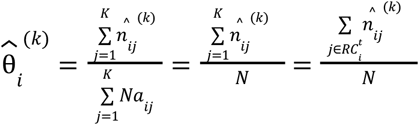

##### 5.4 Zero-count equiRCs

Since long read RNA-Seq does not include a fragmentation step, most equivalent classes that are theoretically possible based on overlapping transcript intervals will not be observed in practice. For most transcripts (with moderate to high expression) the set of possible read classes is equal to the set of observed read classes. However, for transcripts which are very lowly expressed or inactive, the equiRCs representing the full-length transcript may not be observed, which can lead to an overestimation when such transcripts share read classes with highly expressed transcripts. To address this, we define the set of possible read classes as the set of observed read classes plus the set of full length read classes which are not observed. We set the read count for non-observed read classes to zero (zero-count, or empty read classes). This set of read classes will be used for quantification and to obtain the parameter *a_ij_*.

##### 5.5 Full-length and unique support

To keep track of full-length reads, we define a (observed) indicator variable *I^f^_ij_* which takes a value of 1 if reads in read class *j* that originate from transcript *i* are full length reads and 0 otherwise. We can use *1* – *I^f^_ij_* to denote the indicator variable when reads in read class *j* that originate from transcript *i* are partial (non-full-length) reads. We then obtain the full-length transcript estimate for each transcript 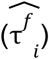 as:

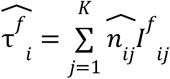

And we have the partial transcript estimate for each transcript 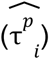 as

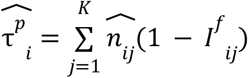

Similarly, to keep track of unique read support, we define a (observed) indicator variable *I^u^_ij_* which takes a value of 1 if reads in read class *j* that originate from transcript *i* are uniquely mapped to transcript *i* and 0 otherwise. We then obtain the unique transcript estimate for each transcript 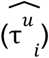 as

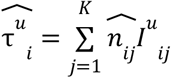

Implementation of Bambu was done in R using Rcpp ^41,42^, Bambu is available through bioconductor.

### Long read RNA-Seq data

For this analysis, we have used SG-NEx core cell lines include A549, K562, Hct116, HepG2, MCF7, HEYA8 and human embryonic stem cell (hESC) cell line generated using cDNA, direct cDNA, and direct RNA protocols. Processed fastq and genome alignment bam files were used for different methods.

### Transcript discovery evaluation

To evaluate the accuracy of the model in identifying valid novel transcript candidates, we split the read classes according to the chromosomes. We used read classes from chromosomes other than the spike-in chromosome and chromosome 1 that overlap with any annotated genes and with at least 2 supporting reads to train the model. The model was trained for each sample separately. Sensitivity is measured as a percentage of the expressed annotated transcripts in the sample (read classes that are assigned as equal). Additionally, to evaluate the robustness of the model, we also trained a default model for when there is not enough data to train on, using a random SG-NEx sample. We then used the two models to predict the transcript probability scores for all read classes with at least 2 supporting reads from the spike-in chromosome and chromosome 1.

To measure the interpretability of the NDR threshold, we ran Bambu without the annotations for chromosome 1 on all SG-NEx samples. We then measured the precision as the fraction of read classes from chromosome 1 that were below the varying NDR thresholds matching the splicing junctions of reference annotations from chromosome 1. This was similarly performed for different read count thresholds. We also compared the precision of Bambu to both StringTie2 and TALON across all SG-NEx samples, which are of varying sequencing depths. To match the default read count threshold used in Bambu, StringTie2 was run with the recommended “-L -G” parameters as well as “-c” at 2 which represents the minimum reads per bp coverage to consider for multi-exon transcript. Similarly, TALON was run withdefault parameters except for --minCount at 2 as part of talon_filter_transcripts which represents the number of minimum occurrences (reads) required for a novel transcript.

To benchmark the performance of Bambu in transcript discovery, we generated a partial annotation where we randomly removed 50% of the annotations on chromosome 1. We then ran transcript discovery with Bambu, FLAIR, StringTie2 and TALON with default parameters where applicable using all SG-NEx core samples individually. For Bambu, we varied the NDR threshold from 0.1 to 1 (0.1, 0.15, 0.2, 0.25, 0.3, 0.35, 0.4, 0.5, 0.6, 0.7, 0.8, 0.9, 1), For StringTie2 we varied the “-c” parameter choosing values between 2 and 50 (2, 4, 6, 8, 10, 15, 20, 30, 50). Due to the higher running time for TALON and FLAIR, we only varied “--minCount” and “-s” in as part of “flair collapse” which represents the minimum number of supporting reads for an isoform from 2 to 10 with a step of 2 respectively. All the novel isoforms were combined with the partial annotations to provide the final output annotation. For each tool, we then evaluated the sensitivity and precision using gffCompare ^43^ with default parameters by comparing the final output annotation with the complete annotations of chromosome 1. gffCompare measured sensitivity as the proportion of transcripts in the reference annotations that were detected. As 50% of the annotations of chromosome 1 are provided, a minimum of 50% is expected. As starts and ends are usually very challenging to determine, we focused on the intron chain level performance which also ignores the presence of single exon transcripts in all tools. To test the performance of these tools across multiple samples, we repeated the above analysis but with all HepG2 samples together. TALON was excluded from this analysis as we could not successfully run the tool using these samples. For StringTie2, we used “--merge” to combine the output annotations from all samples at the different “-c” thresholds described above. StringTie2 was additionally run using “-T” which represents the minimum input transcript-per-million to include in the merge and was performed on the isoforms discovered at “-c 2”.

### Transcript quantification with context-specific annotations

To assess the impact of context-specific annotations on quantification, we generated a partial annotation for the Sequin chromosome^25^ by removing 1 transcript at random for each multiple-isoform gene, and removing single-isoform genes at a random 50% probability. By doing so, we expected that the partial annotation would be missing 40-50% of its transcripts. The partial sequin annotations were then added to the Grch38 Ensembl annotations release version 91. We varied the novel discovery rates (NDRs) from 0 to 1 with a 10% increasing gap and applied it on the samples with Sequins (MixA V2) in the SG-NEx core cell line samples. The estimated transcript expression levels are then compared against the expected transcript expression levels per million, calculated by the relative concentration levels times the expected number of Sequin reads per million (1% x 1 million, where 1% is the spike-in percentage), and the estimates when the Sequin chromosome is provided at full. For the full annotation analysis, we applied Bambu without discovery, i.e., NDR = 0.

We benchmarked Bambu against transcript-discovery assisted quantification approaches, StringTie2, FLAIR, TALON, and quantification-only approaches, LIQA, NanoCount, Salmon and featureCounts. Genome bam files and unmapped fastq files are used as input per condition as described below. For StringTie2, we first performed StringTie2 with the recommended “-L -G” parameters on bam files to discover novel transcripts in each sample. We then performed “ stringtie --merge” to combine novel transcripts across samples. Lastly, we repeated the first step with “-B -e” parameters added and the merged gtf following “ -G” parameter to quantify the transcripts across samples. For FLAIR, we generated bed12 files using bam files with secondary alignments removed, and then performed “flair correct” on those bed12 files to obtain psl files. With psl files, we then performed “flair collapse” on all samples concatenated fastq file to obtain a collapsed fasta file. Lastly, we performed “flair quantify” with the collapsed fasta file and each fastq file again to quantify for each sample. For TALON, we first initialised the database with “talon_inititalize_database” and we then performed “talon_label_reads” for each sample bam file with a MD tag added. We then performed “talon” to discover novel transcripts and after which, “talon_abundance” to quantify transcript expression for each sample. For LIQA, we first filtered bam files with “samtools -F 2308 -q 50” and then performed “liqa -task quantify” with filtered bam files and “-max-distance 10 -f_weight 1” as recommended in the manual. For NanoCount, we used the recommended “-ax map-ont -p 0 -N 10” to align reads to transcriptome reference with minimap2 (version 2.17) ^44^ and output TPM values are used in comparison. Note that for NanoCount, we excluded the top three largest samples as we were unable to run NanoCount successfully due to memory issues. For Salmon, we followed the ONT pipeline (https://github.com/nanoporetech/pipeline-transcriptome-de) where we used “-ax map-ont -p 1.0 -N 100” parameters to align reads to transcriptome reference with minimap2 (version 2.17) and then “quant --noErrorModel -l U” parameters for salmon. For completeness, we have also included FLAMES ^39^ and IsoQuant ^45^, two other available long read transcript discovery and quantification methods for the quantification benchmark. For FLAMES, we processed fastq files following the usage with bulk data analysis suggestion with example SIRV_config.json. For IsoQuant, we processed bam files with “--data-type nanopore” parameter. For StringTie2, FLAIR, IsoQuant and Salmon, the output transcript per million (TPM) expression levels are used for comparison. For TALON, FLAMES, LIQA and featureCounts, the reported counts per transcript were normalised to counts per million (CPM) for comparison. For NanoCount, the output abundance estimates were used for comparison.

### Full-length and unique read support

Bambu provides full-length and unique read support estimation for each transcript in each sample. To assess the impact of estimation uncertainty due to missing data, we performed two analyses.

In the first analysis, we applied CPM, unique read count, and full-length read count based filtering on the SG-NEx core cell lines to assess the influence of filtering in providing more stable estimation. For samples generated using the same protocol within each cell line, we calculated the intra-sample CPM estimates correlation on filtered transcripts based on average CPM, unique read count, or full-length read count passing varying thresholds from 1 to 20, and we took the average correlation.

To evaluate the efficacy of quantification of tools in the presence of in-active transcripts, we generated artificial spliced-isoforms which could be assigned read support without full-length and unique reads. To avoid overly complex scenarios, we selected genes with average expression levels greater than 100 CPM, a partial read support fraction greater than 30%, less than 8 isoforms, and with no isoforms having 3 or more exons. For the selected genes, we identified the most abundant isoform as the reference isoform on which to base the artificial spliced-isoform. From the internal exons of this isoform, we removed the two most commonly used exons among the other isoforms of this gene generating an artificial exon-skipping event. When there are four or more equally commonly used internal exons, we will randomly choose four to remove to mitigate the complexity in such cases. We run Bambu without discovery on the SG-NEx Hct116 samples using the complete annotations that included the artificial transcripts.

To assess the performance of using full-length and unique support to determine if a transcript is truly expressed, we compared CPM, full-length read count, full-length CPM, unique read count, and unique CPM filters at thresholds from 1 to 20. As an additional comparison check, we also ran Salmon and NanoCount using the same artificial annotations, using both SalmonTPM, and NanoCountCPM as filters respectively. For NanoCount, we excluded the two largest samples for this analysis as we were unable to run them with NanoCount due to memory issues. We measured sensitivity as the percentage of transcripts that pass the thresholds for each filtering method among all annotated transcripts, and the precision is calculated as the percentage of valid (non-artificial) transcripts among all transcripts that pass the thresholds for each filtering method. We averaged the sensitivity and precision across the Hct116 samples.

### Quantification of retrotransposon-derived isoforms

To identify retrotransposon-derived genes and isoforms, we focused on the SG-NEx hESC samples and ran Bambu with a NDR threshold of 0.3 as we expected an enrichment of novel transcripts due to the abundance of repeat elements compared to the other SG-NEx cell lines. The extended annotation output from Bambu was then overlapped with the RepeatMasker^46^ sequences matching the Grch38 Ensembl annotations release version 91 ^47^. To identify the top expressed repeat element types, we ranked the number of novel spliced isoforms expressed with a CPM level greater than or equal to that do not overlap with any canonical annotations in any of the hESC samples for each repeat element type.

To quantify the overlapping percentage of repeat elements for each transcript, we looked at the overlap for each exon within each transcript for each repeat. Exons not overlapping with canonical annotations are noted as novel exons. The average overlap percentages were calculated within annotated or novel exons first and then summed up to provide the overall overlap percentage for the transcript. We then compared the distribution of overlapping percentages between annotated isoforms and novel isoforms. We focused on annotated exons in annotated isoforms and novel exons in novel isoforms for this comparison to reduce the potential biasing towards the null hypothesis. A paired t-test^48^ with unequal variance assumption was then conducted to assess the level of significance between the mean overlapping percentages, with two-sided two-sample Kolmogrov-Smirnov test^49^ performed to test the differences in the distributions of the overlapping percentages.

To identify the number of novel genes transcribed from HERVH-LTR7 repeats, we shortlisted novel isoforms, including non-canonical overlapping novel isoforms, re-arranged, or novel spliced canonical isoforms that are overlapping with HERVH-LTR7 repeats, and ranked them by their average expression levels across the hESC samples. We then identified the top expressed isoforms that constitute 90% of the overall isoform expression. To understand the expression of these retro-transposon isoforms in cancer cell lines, we performed Bambu with the extended annotation obtained from hESC samples on cancer cell lines for quantification. Note that for this analysis, HEYA8 samples were not included due to a sample cross-contamination in reads caused by the embedded de-multiplexing protocol used by Guppy (version 3.2.10) ^50^ during the basecalling process. We also performed Bambu on hESC samples without discovery to understand the impact of not discovering these retro-transposon genes/isoforms.

### Benchmark on running time and memory usage

Running time and usage were evaluated with two scenarios. In the first scenario, we evaluated different methods when processing 10 samples with varying sequencing depth (500K to 4.5 million reads) using a single CPU. In the second scenario, we evaluated how different methods perform when processing 10 samples in a combined mode using 5 cpus. In the second scenario, only methods that allow customised multi-threading were included.

## Code Availability

Bambu is a R package for transcript discovery and quantification across multiple samples that is maintained on Bioconductor: https://www.bioconductor.org/packages/bambu/. We used the BambuManuscriptRevision branch version of Bambu for analysis done in this manuscript: https://github.com/GoekeLab/bambu. All analysis code will be made available through code ocean.

## Data availability

The data from the SG-NEx project is available through github (https://github.com/GoekeLab/sg-nex-data) and ENA (PRJEB44348).

## Author contribution statement

Y.C. and A.S. designed and implemented the computational method. J.G. conceived the project. Y.C., A.S., and J.G. designed the study and experiments and analysed data. Y.K.W., K.Y., J.J.X.L., and M.H.L. contributed to implementation of the computational method. M.L. contributed to the design of the computation method. Y.C., A.S. and J.G. organised and wrote the paper with contributions from all authors.

## Acknowledgements

This work was supported by funding from A*STAR. M.L. was supported by R01 HG009937.

---

**Supplementary Figure 1.**
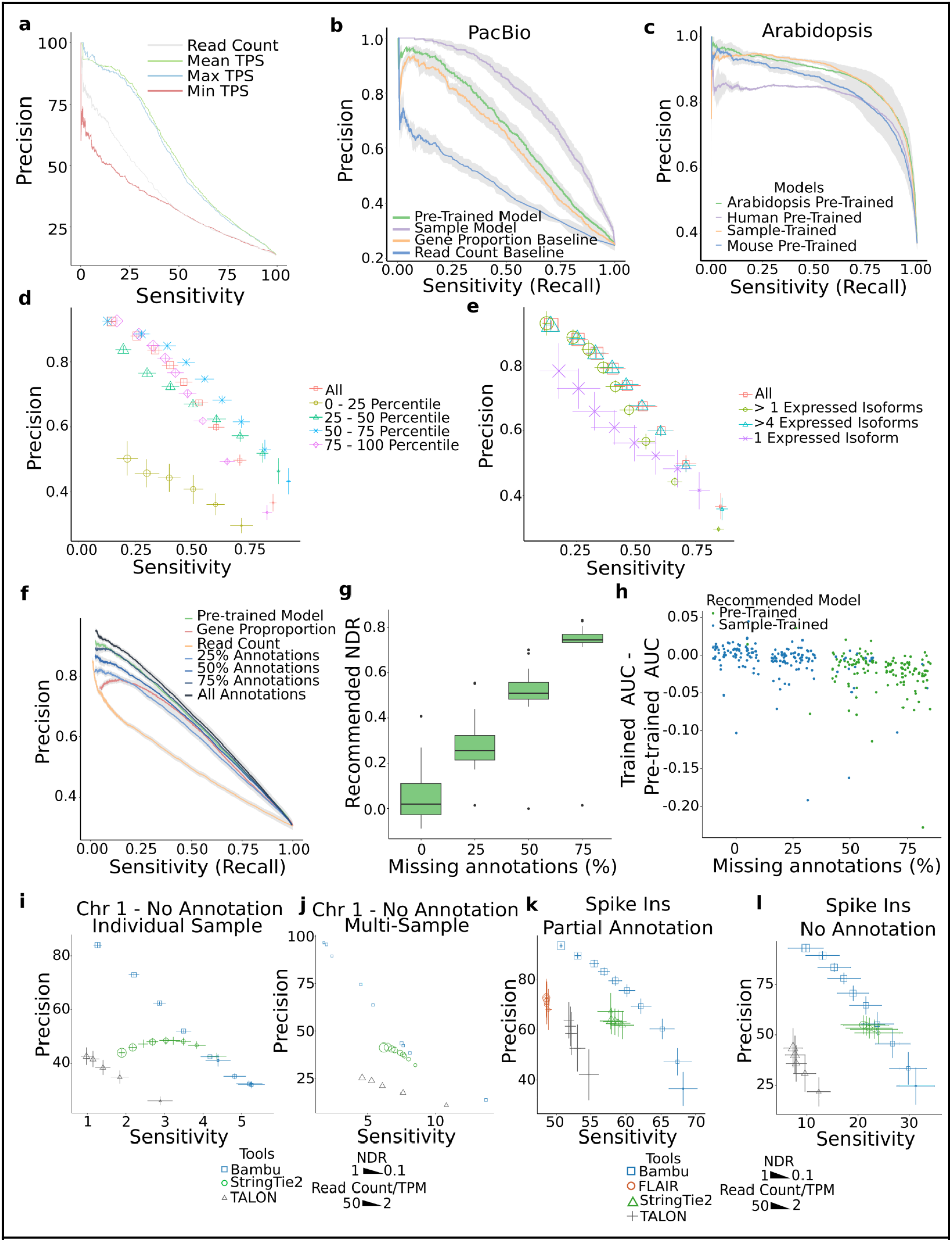
The transcript discovery model is robust and effective at classifying full-length read classes. **(a)** PR curves for the performance of transcript discovery when using minimum, mean, and maximum to combine TPS across samples for the same read class on all HepG2 samples together without annotations for human chromosome 1. The performance of using the sum of read counts as the classifier across the samples is used as a comparison **(b)** A precision recall curve showing the performance of Bambu on PacBio data using either the PacBio trained model (purple), the pretrained model (green), or ranking read classes by gene proportion (orange) or read count (blue). The grey shaded area represents the mean +/- SE of the precision for each line. **(c)** A precision recall curve showing the performance of models pre-trained on human (purple), mouse (blue) or arabidopsis (green) data applied to another arabidopsis tissue. Additionally the performance of the sample trained model is included (orange). The grey shaded area represents the mean +/- SE of the precision for each line. **(d)** The precision and sensitivity of varying Bambu thresholds when looking at a subset of read classes divided into expression quantiles. The same model is applied to all subsets. The full data is coloured in red, read classes that have expressions ranging from 0 to the lower quartile are shaded in yellow, those ranging between the lower quartile and the median in green, between the median and upper quartile in blue and the upper quartile to the max in red. Each subset should represent approximately 25% of the read classes. The lowest expressed quartile is larger than the others due to a greater than 25% of read classes sharing a read count of 2. **(e)** The precision and sensitivity of varying Bambu thresholds when looking at a subset of read classes divided by the number of expressed isoforms (> 2 read count) their gene contains. The same model is applied to all subsets. The full data is coloured in red, the subset of read classes that are the only expressed isoforms in their gene are coloured purple, those which have two or more isoforms are shaded in teal, and those with five or more isoforms are green. **(f)** Precision and Recall curves of the classification using Bambu models trained using reference annotations missing a random fraction of annotations used from the human reference annotations (excluding chromosome 1). The models trained using these annotations, are used to classify read classes from chromosome 1. The Pretrained Model represents the in-built model in Bambu which is used when the annotations do not support training and Read Count classifies the read classes solely using read count alone. These were applied to all SG-NEx datasets. **(g)** We measured the difference in ROC AUC of trained and pretrained models in which the trained model was trained with missing reference annotations. Samples in which Bambu recommends using the pretrained model are coloured red (the recommended NDR was calculated as > 0.5), and samples where the trained model is used are coloured blue. **(h)** A box plot showing the distribution of NDR recommendations of SG-NEx samples when Bambu was run with different percentages of reference annotations. **(i)** The average sensitivity and precision of transcript discovery on core SG-NEx samples without annotations for human chromosome 1. Each tool is displayed at several different parameter thresholds: Bambu (blue) with NDR thresholds varying between 1 and 0.1, FLAIR (red), StringTie2 (green) and TALON (greygray) with read count/coverage thresholds varying between 2 and 10, with 4 additional thresholds for StringTie2 at 15, 20, 30 and 50. Horizontal error bars represent the mean +/- SD of the sensitivity and vertical error bars represent the mean +/- SD of the precision **(j)** The measured sensitivity and precision of transcript discovery when combining HepG2 samples, without annotations for human chromosome 1. Each tool is displayed at several different parameter thresholds: Bambu (blue) with NDR thresholds varied between 1 and 0.1, StringTie2 (green) and TALON (grey) with read count/coverage thresholds varying between 2 and 10, and 4 additional thresholds for StringTie2 at 15, 20, 30 and 50 **(k)** The average sensitivity and precision of transcript discovery on spike-in data without annotations for the spike-in chromosome. Each tool is displayed at several different parameter thresholds: Bambu (blue) with NDRthresholds varying between 1 and 0.1, StringTie2 (green), FLAIR (red) and TALON (grey) with read count/coverage thresholds varied between 2 and 10. Horizontal error bars represent the mean +/- SD of the sensitivity and vertical error bars represent the mean +/- SD of the precision **(l)** The average sensitivity and precision from transcript discovery outputs run on SG-NEx data with 50% of the spike-in annotations randomly removed. Each tool is displayed at several different parameter thresholds. Bambu (Blue) was run using novel discovery thresholds between 1 and 0.1. StringTie2 (green), FLAIR (red) and TALON (grey) were run with read count/coverage thresholds between 2 and 10. Error bars represent the standard error

**Supplementary Figure 2.**
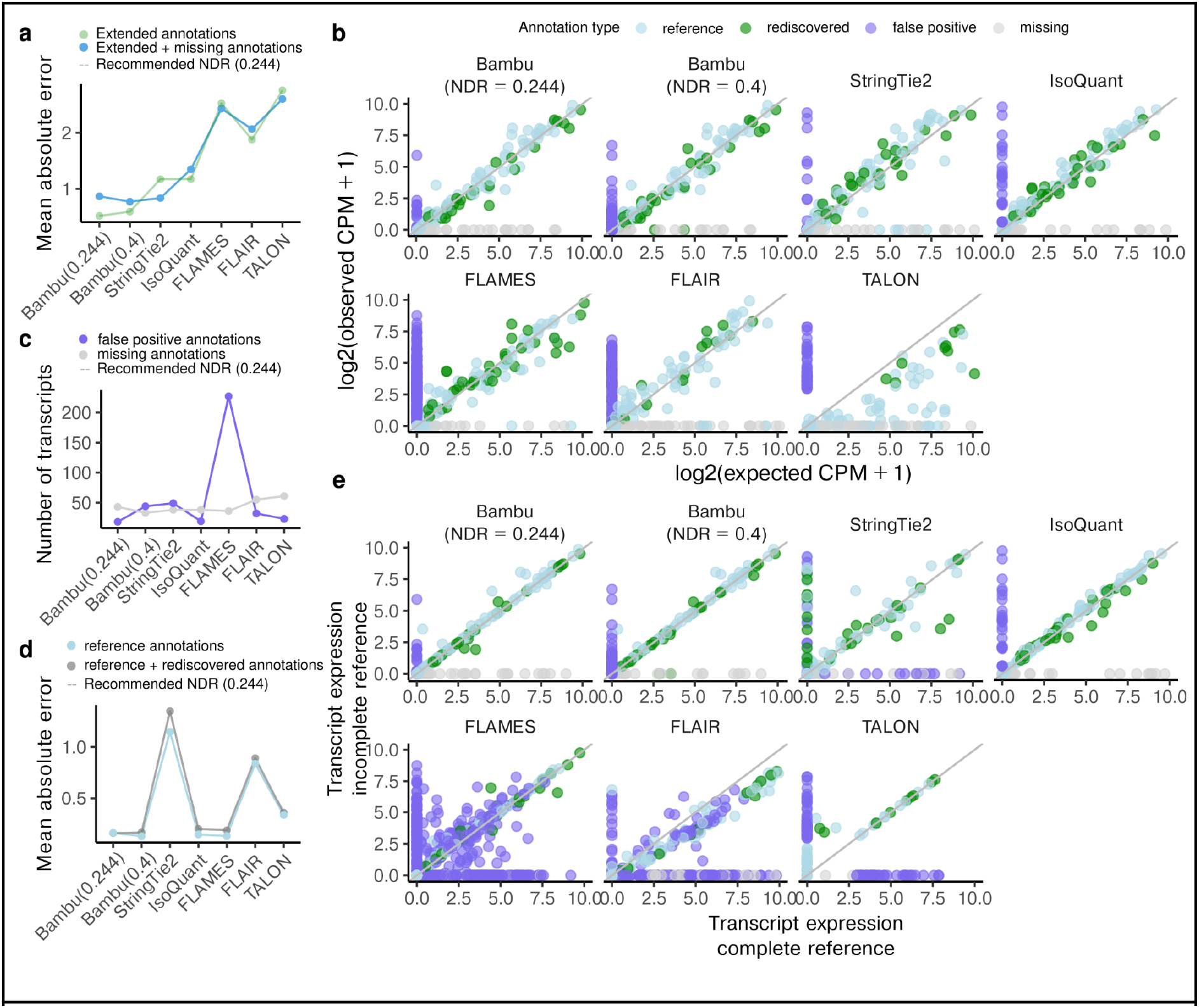
Quantification accuracy and consistency by methods that does both transcript discovery and quantification. **(a)** The mean absolute error between the log2 normalised spike-in transcript abundance estimates and log2 normalised expected spike-in abundance when applying Bambu with recommended NDR (0.244), Bambu with NDR = 0.4, StringTie2, IsoQuant, FLAMES, FLAIR, and TALON for annotations for extended annotations by each of these methods, including annotations that are present in the reference (partial) sequin annotations, the annotations that have been artificially removed and rediscovered by each of the methods, and also the false positive annotations discovered by each method (green), plus annotations that have been artificially removed from the partial annotation and remained missing after transcript discovery, i.e., missing annotations (blue) **(b)** Scatterplots between log2 normalised transcript abundance estimates and log2 normalised expected spike-in abundance when applying Bambu with recommended NDR (0.244), Bambu with NDR = 0.4, StringTie2, IsoQuant, FLAMES, FLAIR, and TALON for spike-in transcripts annotated transcripts (light blue), transcripts artificially removed from the reference and rediscovered by Bambu (green), false positive transcripts (purple), transcripts artificially removed from the reference and remained missing after Bambu discovery (grey) **(c)**The number of missing (grey) and false positive (purple) transcripts using partial full sequin annotations when applying Bambu with default recommended NDR (0.244), Bambu with NDR = 0.4, StringTie2, IsoQuant, FLAMES, FLAIR, and TALON. **(d)** and full annotations when applying Bambu with default NDR, Bambu with NDR = 0.4, StringTie2, IsoQuant, FLAMES, FLAIR, and TALON. This is calculated separately for annotated transcripts (light blue), transcripts artificially removed from the reference (green), and false positive transcripts (purple) **(e)** Scatterplots between log2 normalised transcript abundance estimates with complete annotations and log2 normalised transcript abundance estimates with partial annotations when applying Bambu with default NDR, Bambu with NDR = 0.4, StringTie2, IsoQuant, FLAMES, FLAIR, and TALON for spike-in transcripts annotated transcripts (light blue), transcripts artificially removed from the reference and rediscovered by Bambu (green), false positive transcripts (purple), transcripts artificially removed from the reference and remained missing after Bambu discovery (grey)

**Supplementary Figure 3.**
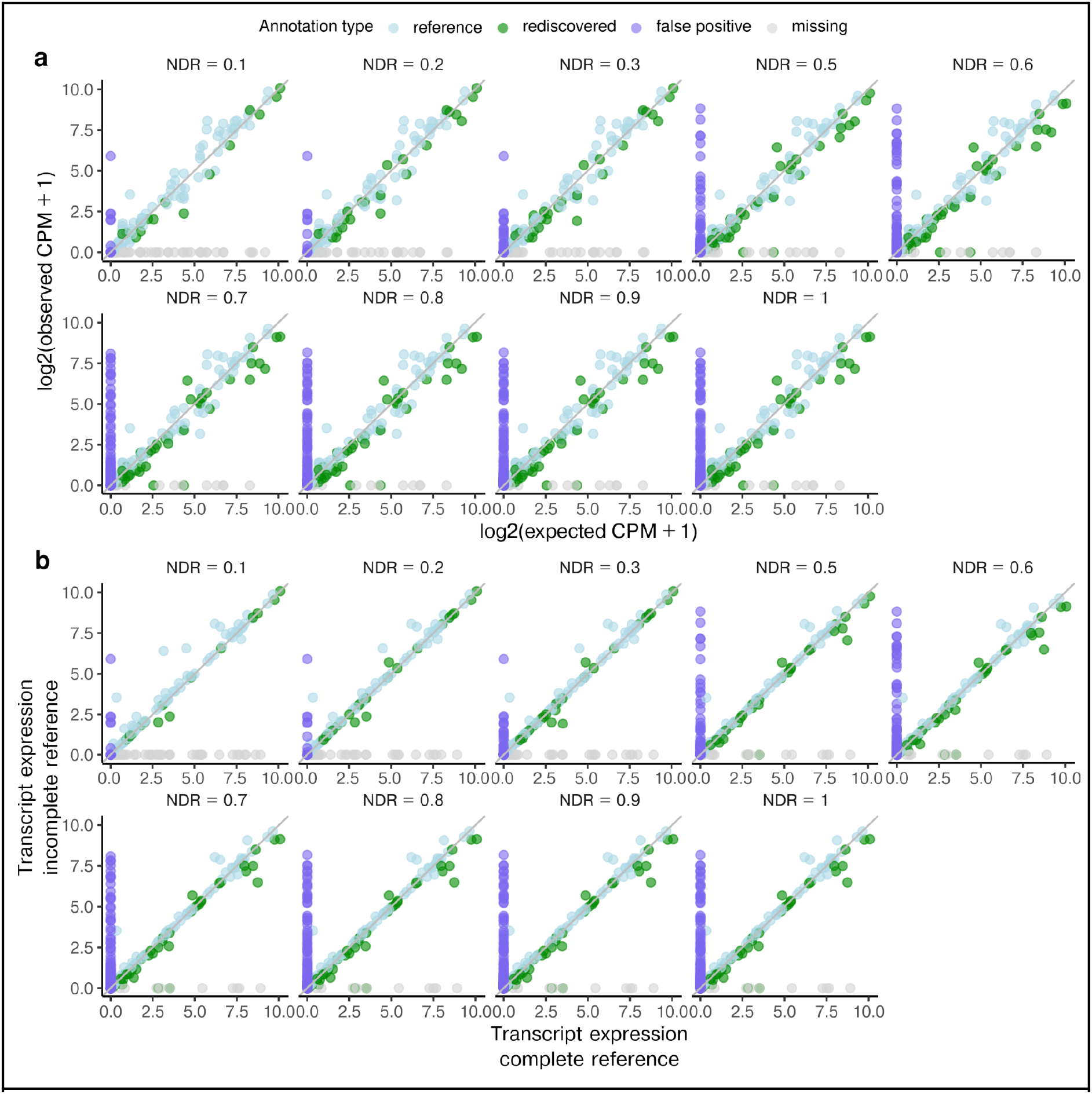
Comparison of quantification for Bambu with other NDRs. **(a)** Scatterplots between transcript abundance estimates and expected spike-in concentrations for spike-in annotated transcripts (light blue), transcripts artificially removed from the reference and rediscovered by Bambu (green), false positive transcripts (purple), transcripts artificially removed from the reference and remained missing after Bambu discovery (grey) when applying Bambu with varying NDRs from 0.1 to 0.3, and 0.5 to 1**(b)** Scatterplots between log2 normalised transcript abundance estimates with complete annotations and log2 normalised transcript abundance estimates with incomplete annotations for spike-in transcripts annotated transcripts (light blue), transcripts artificially removed from the reference and rediscovered by Bambu (green), false positive transcripts (purple), transcripts artificially removed from the reference and remained missing after Bambu discovery (grey) when applying Bambu with varying NDRs from 0.1 to 0.3, and 0.5 to 1

**Supplementary Figure 4.**
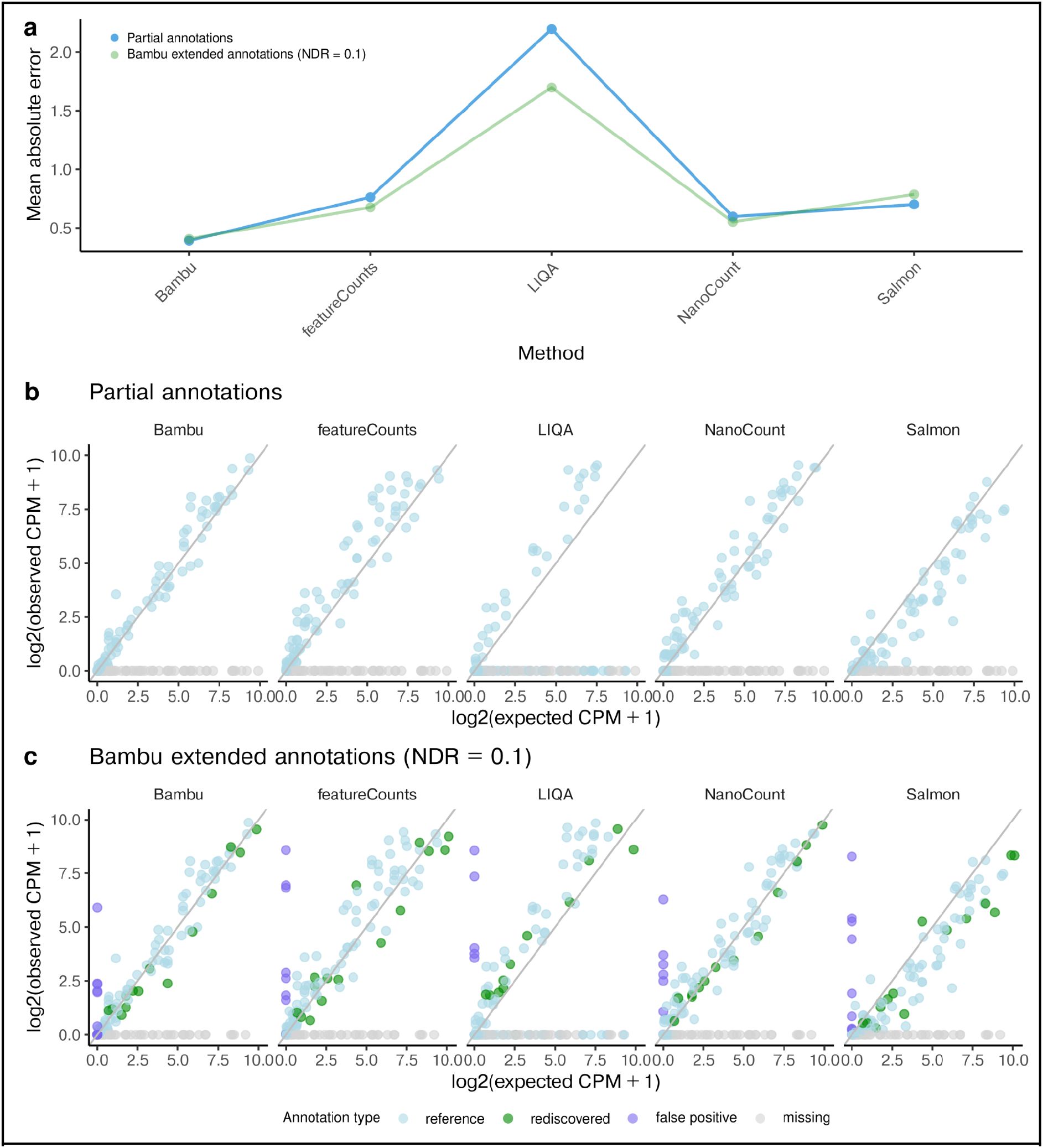
Quantification performance when partial/extended annotations are provided. **(a)** The mean absolute error between the log2 normalised spike-in transcripts estimates and log2 normalised true concentration levels when applying Bambu, featureCounts, LIQA, NanoCount, and Salmon with partial (blue) or Bambu extended (NDR = 0.1, green) annotations for transcripts that are present in the reference (partial sequin annotations). **(b-c)** Scatterplots between transcript abundance estimates and expected spike-in estimates when applying Bambu, featureCounts, LIQA, NanoCount, and Salmon for transcripts that are present in the reference (partial sequin annotations) (light blue), transcripts that have been artificially removed from the reference and rediscovered by Bambu (green), false positive transcripts (purple), transcripts that have been artificially removed from the reference and remained missing after Bambu discovery (grey): (b) with partial annotations; (c) with extended annotations by Bambu (NDR = 0.1).

**Supplementary Figure 5.**
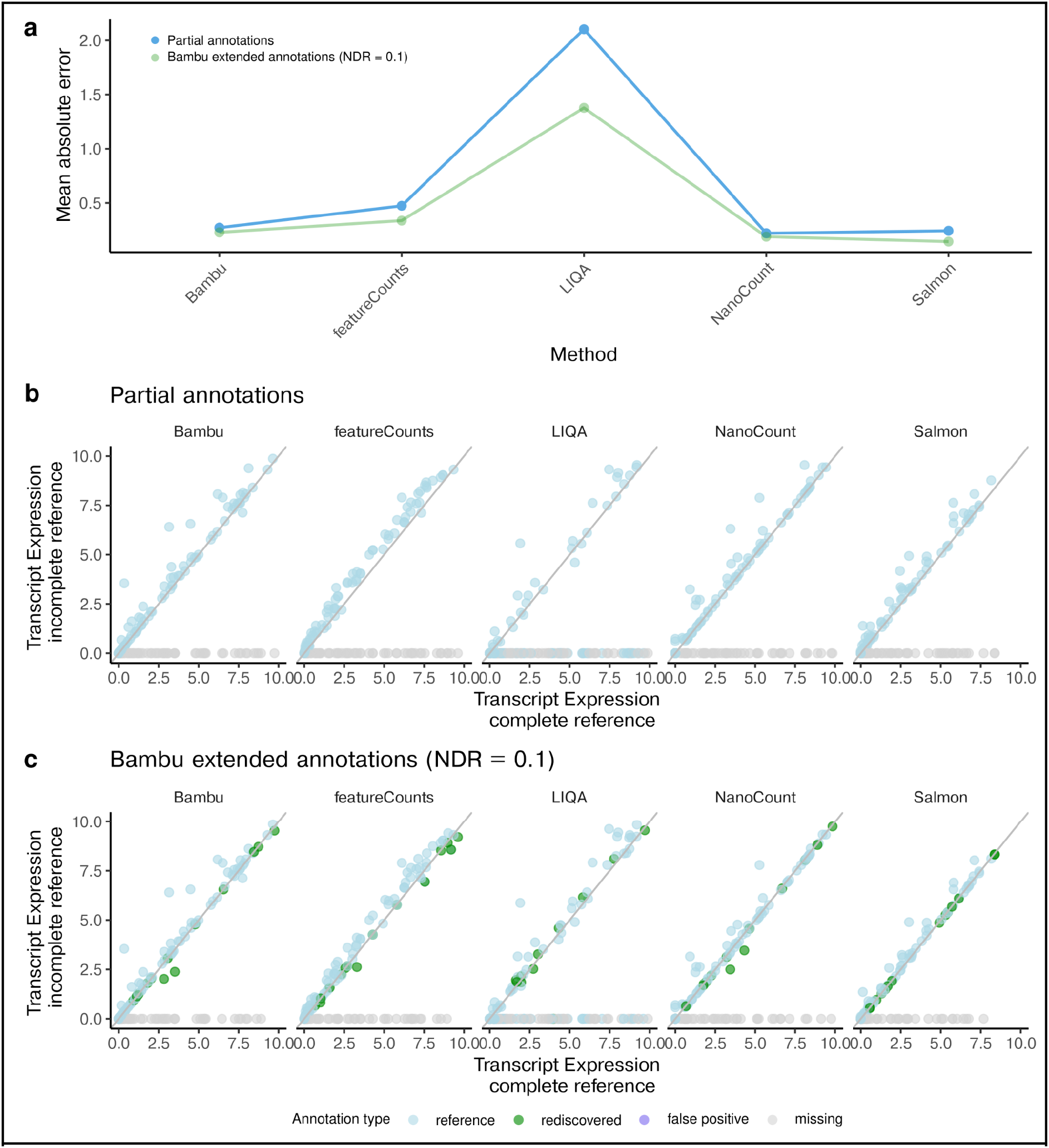
Quantification when partial/extended annotations are provided compared against quantification when complete annotations are provided. **(a)** The mean absolute error between the log2 normalised spike-in transcript abundance estimates with partial (blue) or Bambu extended (NDR = 0.1, green) annotations against with full annotations when applying Bambu, featureCounts, LIQA, NanoCount, and Salmon for transcripts that are present in the reference (partial sequin annotations). **(b-c)** Scatterplots between log2 normalised transcript abundance estimates with complete annotations and log2 normalised transcript abundance estimates when applying Bambu, featureCounts, LIQA, NanoCount, and Salmon for transcripts that are present in the reference (partial sequin annotations) (light blue), transcripts that have been artificially removed from the reference and rediscovered by Bambu (green), false positive transcripts (purple), transcripts that have been artificially removed from the reference and remained missing after Bambu discovery (grey): (b) with partial annotations; (c) with extended annotations by Bambu (NDR = 0.1).

**Supplementary Figure 6.**
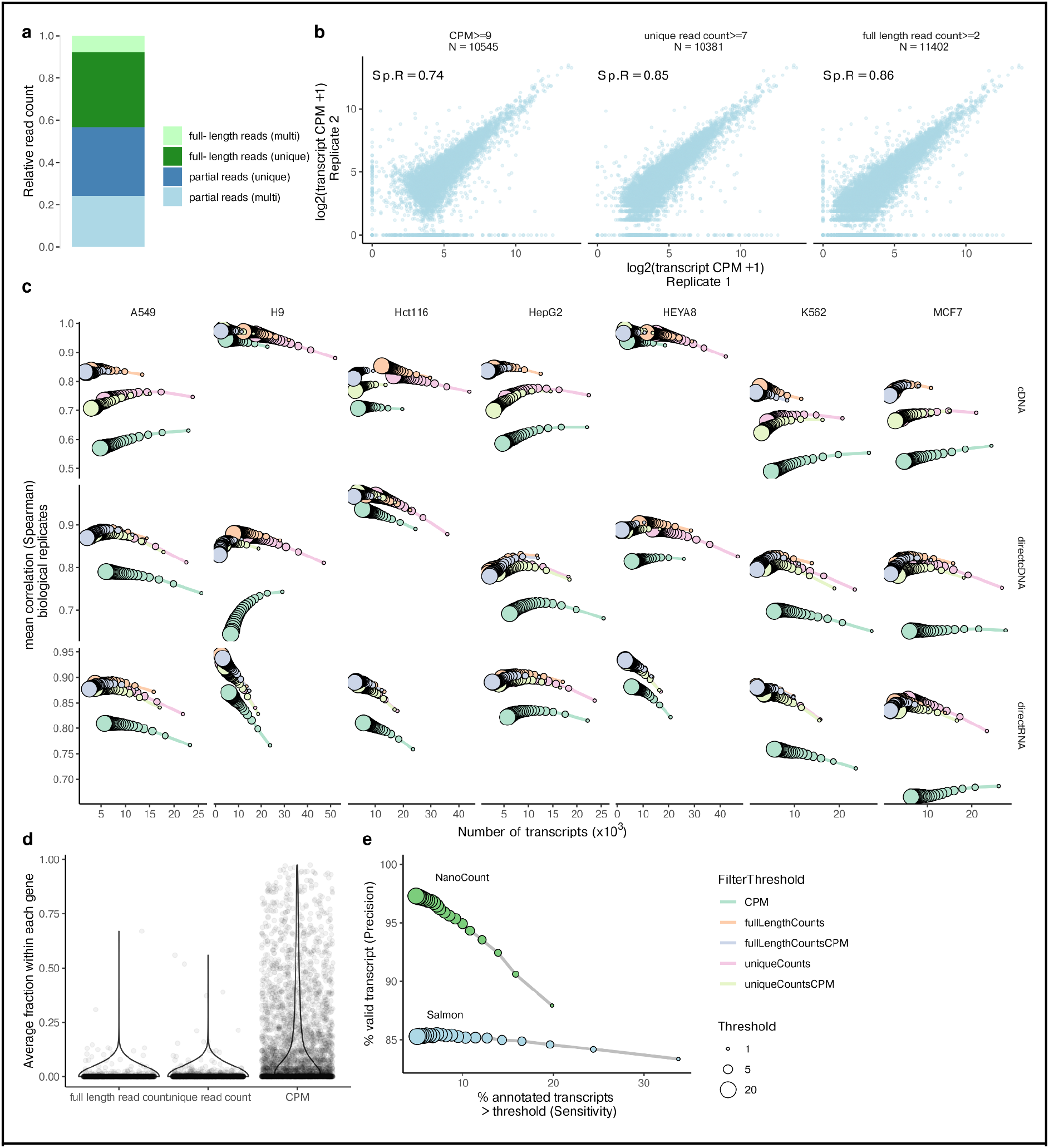
Full-length and unique read support show evidence additional to transcript abundance estimates. **(a)** The distribution of average reads categorised as full-length, partial length, and unique **(b)** The comparison of transcript abundance estimates between two replicates of MCF7 generated using the direct cDNA protocol, with filtered transcript number being approximately 30 thousand, when filtered using CPM, unique read count, and full-length read count. The spearman correlation is shown in the top right for each filter **(c)** The mean spearman correlation of transcript abundance estimates between replicates for each cell line generated using cDNA, direct cDNA, and direct RNA protocols against number of transcripts that pass the filter, using CPM, full-length read count, full-length CPM, unique read count and unique CPM filter thresholds, with thresholds ranging from 1 to 20 **(d)** Violin plot showing the median, upper and lower quartiles, and 1.5 x interquartile range of the average fraction of full-length, unique read counts and CPM for each artificial transcript across Hct116 samples **(e)** The sensitivity and precision of NanoCount and Salmon abundance estimates in filtering transcripts overlapping with highly abundant isoforms that have no unique or full-length reads support at varying filtering thresholds from 1 to 20 on Hct116 samples. Filtering was based on the average values across replicates being not lower than the threshold.

**Supplementary Figure 7.**
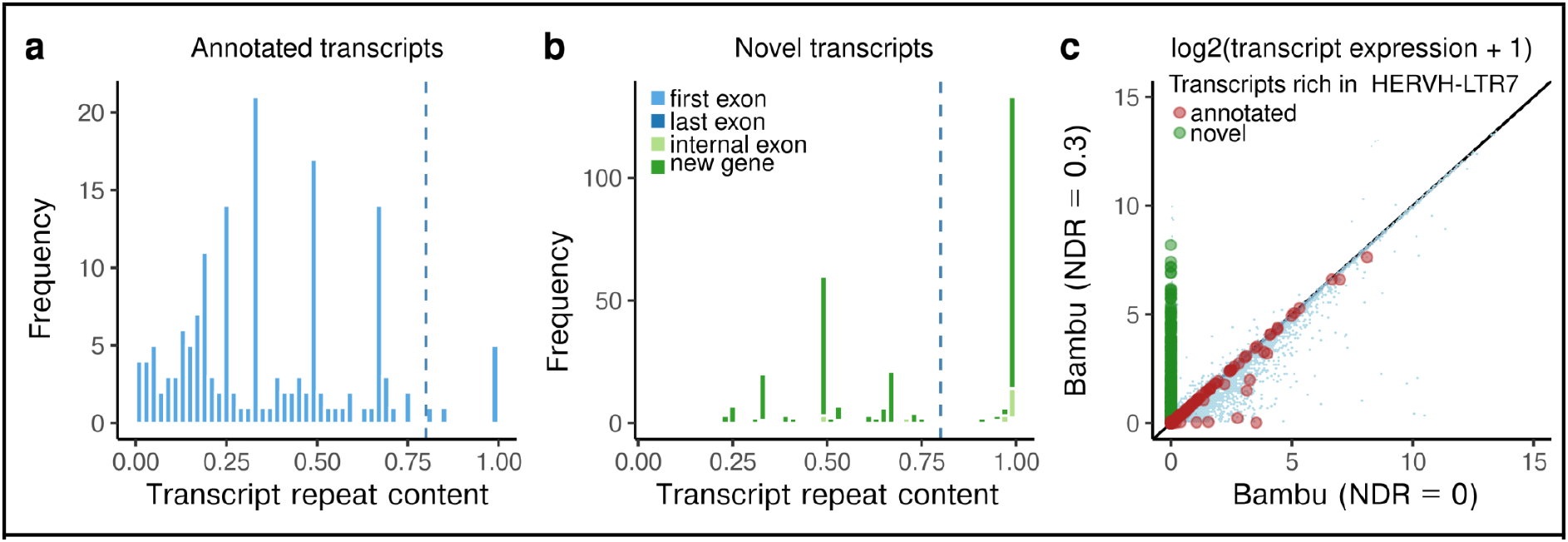
Transcript discovery identifies novel transcripts overlapping highly with repeats. **(a-b)** Histogram of overlapping percentage of repeats for (a) annotated transcripts and (b) novel transcripts **(c)** Scatterplot of transcript abundance estimates with discovery (NDR = 0.3) against that without discovery (NDR = 0), with red and green points showing annotated and novel transcripts with at least 80% overlapping with HERVH-LTR7, and light blue points showing other transcripts with less than 80% overlapping with HERVH-LTR7

**Supplementary Figure 8.**
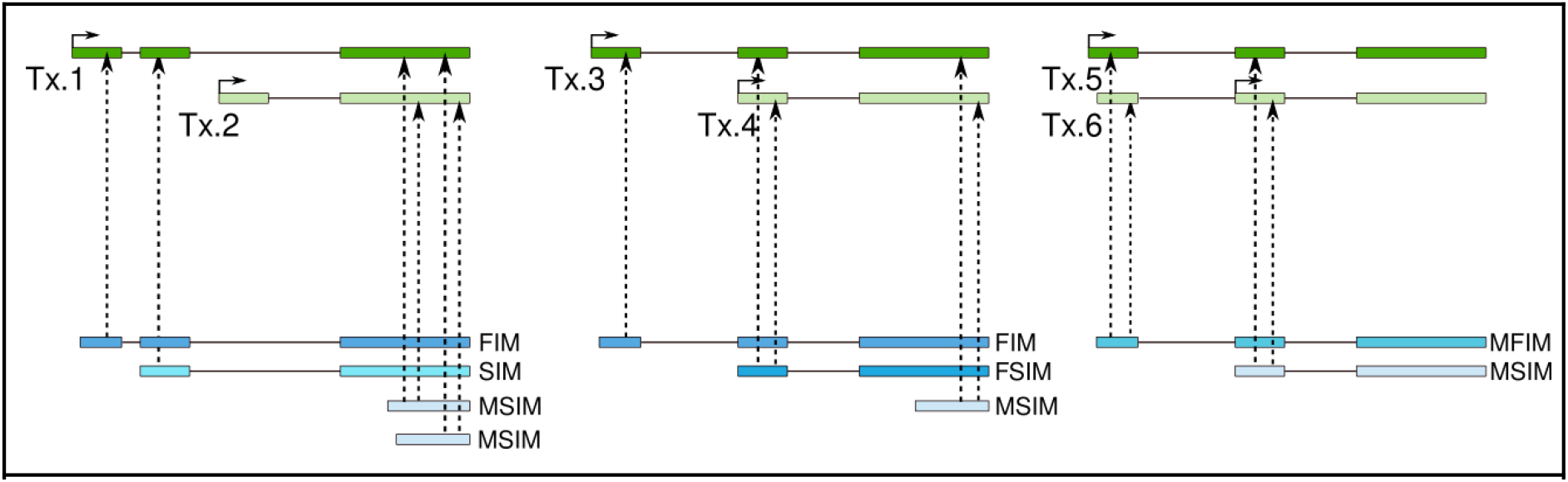
Equivalence Read Class (EquiRC) types. Illustration of the five equiRC types: FIM (Full Intron Match), equally aligning to unique transcript; SIM (Subset Intron Match), partially aligning to unique transcript; FSIM (Full and Subset Intron Match), equally aligning to subset transcripts, while partially aligning to longer transcripts; MFIM (Multiple Full Intron Match), equally aligning to multiple transcripts, usually very similar transcripts; MSIM (Multiple Subset Intron Match), partially aligning to multiple transcripts (these are the mostly fragmented reads)

