## Supplemental Text for "Context-Aware Transcript Quantification from Long Read RNA-Seq data with Bambu"

Affiliations:
1.Genome Institute of Singapore, Singapore, Singapore
2.Yong Loo Lin School of Medicine, National University of Singapore, Singapore
3.Department of Biostatistics, University of North Carolina-Chapel Hill, Chapel Hill, USA
4.Department of Genetics, University of North Carolina-Chapel Hill, Chapel Hill, USA
5.National Cancer Centre Singapore, Singapore, Singapore

*contributed equally

### Supplementary Text

#### Content

**Features used for transcript discovery** 1
**Contribution of transcript features to transcript discovery** 3
**Combining Samples** 4
**Single Exon Read Classes** 4
**Using a pre-trained model (in practice)** 5
**Bambu performance at different levels of annotation completeness** 6
**Filtering incompatible read classes for transcript quantification** 7
**Impact of Minimap2 alignment parameters used on NanoCount and Salmon quantification results** 8
**Running time and memory usage of Bambu against other methods** 10
**Feature comparison** 13

#### 1. Features used for transcript discovery

Bambu's transcript discovery model uses nine features during classification: number of reads, gene proportion, the standard deviation of the starts and ends of read classes, and the number of polyAs and polyTs found at the read class start and ends, which are detailed below:

**Feature definition:**

Each read class i is described by a vector $x_i \in R^9$ :

##### 1. 1 Number of reads ($x_i^{RC}$)

The number of reads for each read class is defined as the number of aligned reads that share the exact exon-junctions ($c_i$), normalised by library size for sample j ($l_j$):

$$x_i^{RC} = \frac{c_i}{l_j}$$

With $l_j = \sum_{i=1}^{M} c_i$.

This feature is an intuitive measure for the validity of a read class and its abundance in the sample of interest, as the read count directly reflects the number of observations for this read class in the data set.

*Limitations of read count as a standalone parameter*

Systematic errors during library preparation, sequencing, or alignment can result in larger read counts for read classes that are not valid transcripts, such as degraded RNAs and non-full-length reads. Furthermore, read classes with low read count can still be valid transcripts, and highly expressed genes are more likely to lead to read classes with high read count that originate from degradation products or other possible artefacts. Therefore, higher read count does not guarantee a high probability that a read class is a valid transcript, and a low read count does not guarantee that a read class is not a valid transcript.

##### 1.2 Gene proportion ($x_i^{GP}$)

Gene proportion represents the proportion of reads that are assigned to one read class amongst all the reads assigned to the same gene:

$$x_i^{GP} = \frac{c_i}{\sum_{t \in I} c_t}$$

With I representing the set of all read classes that overlap the same gene as read class i

Gene proportion directly addresses some of the main limitations of read count ($x^{RC}$) as it reflects the number of reads assigned to each read class relative to the number of reads assigned to the gene.

*Limitations of read count as a standalone parameter*

There are 2 main limitations for gene proportion. Firstly, this feature is strongly influenced by the number of isoforms for each gene and therefore not comparable across different genes. In particular, gene proportion is always 1 for single transcript genes, and much smaller with larger numbers of expressed transcripts. Secondly, the estimation of gene proportion can be inaccurate with low read counts. With a gene read count of 10, any read class with 2 reads will have a gene proportion of 20%, which is already higher than the gene proportion observed for many valid, annotated transcripts from genes with many isoforms.

**1.3 TSS/TES standard deviation ($x_i^{\sigma TSS}$ and $x_i^{\sigma TES}$)**

This feature is calculated by measuring the standard deviation of the locations of the start and end coordinates for all the reads comprising a read class. Differences in the standard deviation may indicate specific properties of valid transcripts that distinguish them from degradation artefacts and other invalid transcript candidates.

**1.4 Strand bias ($x_i^{strand}$)**

The strand bias is calculated as the proportion of reads mapping to the strand with higher read count.

In cDNA samples sequencing can start at both the 5' and 3' end of the transcript (whereas in RNA samples, sequencing always starts at the 3' end). For cDNA samples, a deviation from the average strand bias could represent cases where sequencing did not process correctly in one direction resulting in a systematic early truncation.

**1.5 Start and End polyA/T frequency ($x_i^{start-A}, x_i^{start-T}, x_i^{end-A}, x_i^{end-T}$)**

This feature counts the number of A's (orT's respectively) within the first (or last respectively) 10bp of the start and end of the read class. This feature is based on the genome sequence, not on the read or transcript sequence, and does not represent the presence of polyA tails. The presence of polyA/polyT sequences can result in early truncation of the read. One example is strand invasion that can occur during reverse transcription leading to truncated reads with an abundance of A's at the 5' end. This feature is designed to capture sequencing artefacts due to the presence of A or T rich sequences.

#### 2. Contribution of transcript features to transcript discovery

Bambu trains a model that automatically optimises the contribution of all features for transcript discovery based on the sample and technology provided by the user. On their own, only read counts and gene proportion (and to a lesser amount the standard deviation of both ends) are useful as classifiers (Supplementary Text Figure 1a). However as Bambu can learn non-linear relationships, these features still contribute to the overall classification accuracy, with varying impact depending on the sample. In the sample used for the default pre-trained model, gene proportion is the most important feature, whilst the presence of polyA and polyTs at the start of the read class and the read count showed similar importance (Supplementary Text Figure 1b). As this sample is direct RNA and is therefore stranded, the strand bias feature has no relevance in the model, showing how training is able to adapt to the sample as needed. The relevance of the multi-feature approach is highlighted when looking at the change in contribution of the features across multiple samples and library preparations (Supplementary Text Figure 1c). For example the number of T's and A's at the start of a transcript have more impact on the direct RNA-seq data, reflecting specific characteristics of the direct RNA-Seq protocol.

The integration of features using a supervised machine learning model allows for more dynamic and context-specific transcript discovery. Besides being robust and accurate, this approach also reduces the complexity of threshold calibration from nine potentially relevant features to a single, interpretable probability score.

Bambu also allows users to specify read count and gene read proportion thresholds which can be used instead of the NDR, however, this is not recommended.

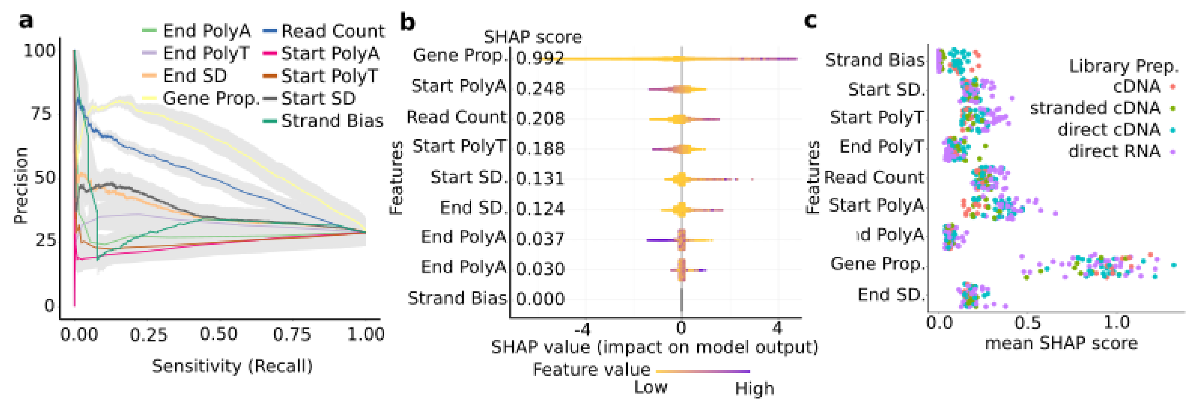


***Supplementary Text Figure 1. Contribution of features to Bambu's transcript discovery model***
**(a)** A precision-recall plot for each feature used in the transcript prediction model in bambu. Features are used to rank read classes/transcript candidates without any transformation. Each precision-recall curve is averaged across all SG-NEx data. The grey shaded area represents the standard error for each feature. **(b)** A SHAP plot showing the value of different features for a model trained on the HepG2 direct RNA replicate 5 run 1 sample. The y-axis is each feature used in the model with the number representing the feature SHAP score (the contribution of the feature) to the model. Each point represents one read class and the colour represents if the feature has a higher value (purple) relative to the feature. The x-axis is the importance of the features value to the model prediction score. **(c)** A jitter plot of the feature SHAP scores for all SG-NEx samples. Each feature from the model is on the y-axis and the mean SHAP score is shown on the x-axis. Each point is coloured based on the library preparation of the sample: PCR cDNA (red), PCR cDNA stranded (green), direct cDNA (blue) and direct RNA (purple)

#### 3. Combining Samples

Bambu trains a model on each sample individually and thereby assigns a different TPS to a read class that occurs across multiple samples. Bambu uses the maximum TPS to combine multiple samples (see Methods for details). As a comparison, we also measured the predictive power with a PR curve when either the maximum, minimum or mean is used to integrate the TPS across samples. Using the mean or max TPS resulted in the best and very similar performance with taking the minimum TPS performing worse than the read count baseline (Supplementary Figure 1a).

#### 4. Single Exon Read Classes

Overlapping single exon reads are combined into single exon read classes. By default Bambu does not report single exon novel transcripts, however users can choose to include them if desired using advanced parameters (see online documentation).

During model training, bambu separately trains single-exon and multi-exon read classes and produces two distinct models. The read classes are not trained together as the features between the two read class types have differing behaviours which negatively affect model performance. Single exon read classes which are wholly contained within an annotated single exon annotation are considered as equal for training purposes (irrespective of how much the start and end sites differ). The TPS for multi-exon and single-exon read classes are predicted by their respectively trained models.

Single exon transcripts can be a source for false positives. While these transcripts can be included, we recommend that they are carefully inspected and interpreted. Single exon read classes are always used for quantification regardless of how they are handled during transcript discovery.

We have performed the quantification by other tools after removing the single exon transcripts. Compared to no filtering on novel single exon transcripts , the number of false novel transcripts is reduced for StringTie2, FLAIR, and TALON, reducing the mean absolute error (Supplementary Text Figure 2, other methods did not report single exon transcripts for the sequin genes).

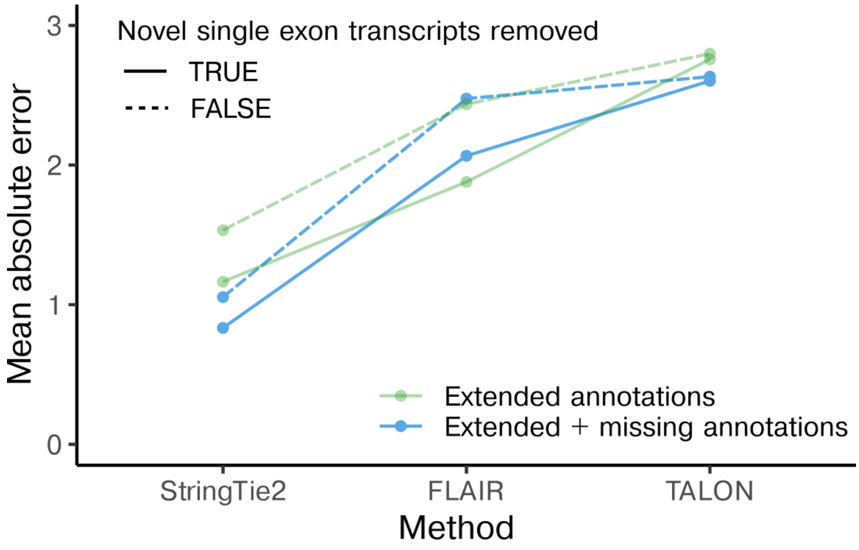


***Supplementary Text Figure 2. Impact of novel single exon transcripts on quantification accuracy***
The mean absolute error between the log2 normalised spike-in transcripts estimates and log2 normalised true concentration levels when applying StringTie2, FLAIR, and TALON with (solid line) or without (dashed line) novel single exon transcripts removed for software annotations, i.e., partial annotations + positive annotations from each of the methods (green), or complete annotations, i.e., full reference annotations + positive annotations for each of the methods (blue).

#### 5. Using a pre-trained model (in practice)

The pretrained model is shown to be able to effectively classify novel transcripts when tested on data from different technology (PacBio) and on different organisms (Arabidopsis) with only a small performance drop compared to a sample trained model (Supplementary Figure 1b). However, for users that do not have sufficient annotations to train their sample, and would like to increase the performance of transcript discovery, Bambu allows for training new models using related data for which comprehensive reference annotations are available. To evaluate this functionality, we trained Bambu on mouse, human and Arabidopsis data, and tested the pre-trained models on a different genotype of Arabidopsis. Here we observe that both the human and mouse pre-trained models can be used to rank novel transcripts in Arabidopsis (Supplementary Figure 1c). However, the model that was trained on *A. Thaliana* and applied to another genotype of *A. Thaliana* showed higher performance that was comparable with the sample/genotype specific model (Supplementary Figure 1c). These results suggest that the generic pre-trained model is robust and can be used effectively, while a pre-trained model that matches the species of interest is expected to lead to an improved performance.

Please refer to the online documentation (https://github.com/GoekeLab/bambu/tree/BambuManuscriptRevision) for details on model training.

#### 6. Bambu performance at different levels of annotation completeness

| Reference annotations | Mean number of expressed annotated transcripts | Mean Model performance. (PR AUC) | Mean ROC AUC |
|---|---|---|---|
| 25% | 2345 | 0.597 | 0.770 |
| 50% | 4688 | 0.639 | 0.785 |
| 75% | 6999 | 0.663 | 0.792 |
| 100% | 9260 | 0.677 | 0.798 |
| Pretrained Model | NA | 0.668 | 0.792 |
| Read Count | NA | 0.498 | 0.678 |

***Supplementary Text Table 1. Performance of Transcript Discovery Model trained with missing reference annotations***
Reference annotations represent the random fraction of annotations used from the human reference annotations (excluding chromosome 1). The models trained using these annotations, are used to classify read classes from chromosome 1. The Pretrained Model represents the in-built model in Bambu which is used when the annotations do not support training and included chromosome 1 during training. Read Count classifies the read classes solely using read count alone. These were applied to all SG-NEx datasets.

| Missing Annotations | 0% | 25% | 50% | 75% | Number of Samples |
|---|---|---|---|---|---|
| PacBio Human | 0.141 ±0.004 | 0.349 ±0.009 | 0.569 ±0.001 | 0.766 ±0.006 | 3 |
| Mouse | 0.128 ±0.010 | 0.336 ±0.010 | 0.557 ±0.015 | 0.774 ±0.004 | 4 |
| Arabidopsis | 0.103 ±0.007 | 0.309 ±0.009 | 0.527 ±0.002 | 0.755 ± 0.002 | 3 |

***Supplementary Text Table 2. NDR recommendation on different datasets***
The table shows the mean recommended NDR across the samples tested. ± is the standard deviation of the mean

#### 7. Filtering incompatible read classes for transcript quantification

To improve Bambu quantification accuracy, we only assign reads to transcripts if they are below a maximum alignment distance of 35bps (referred to as *compatible*), whereas any other read will be excluded from transcript quantification (*incompatible* reads). The filtering step provides more accurate transcript quantification (Supplementary Text Figure 3a-b).

Most reads which are incompatible with transcripts due to missing annotations can still be accurately assigned to genes. Usually, gene expression is estimated as the sum of transcript expression, therefore, when incompatible reads are removed, gene expression will be underestimated. To prevent this, Bambu assigns all incompatible reads to an artificial “unidentified transcript” that is associated with each gene. Gene expression is then estimated using all reads that can be assigned to the transcripts of each gene, including reads that are incompatible with all existing annotations, leading to improved gene expression quantification (Supplementary Text Figure 3c-d).

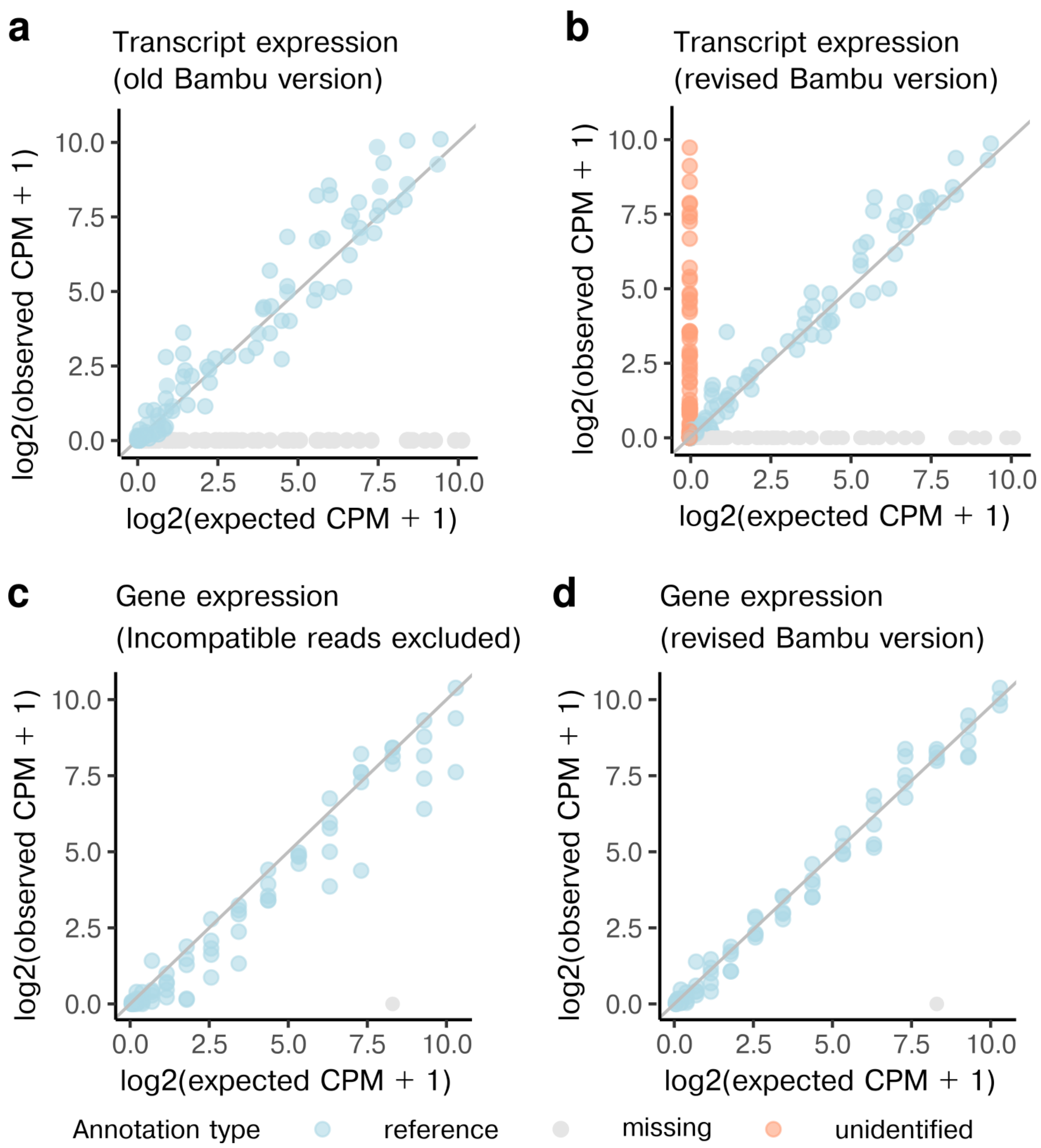


***Supplementary Text Figure 3. Tracking of incompatible reads improves gene expression quantification***

Shown is the scatter plot of observed CPM vs expected CPM for sequin transcripts and genes when Bambu is used with NDR = 0 and partial annotation is provided: **(a)** sequin transcript expression (old Bambu version) vs **(b)** sequin transcript expression (revised Bambu version); **(c)** sequin gene expression when incompatible reads are excluded vs **(d)** sequin gene expression when incompatible reads are included (revised version of Bambu). Blue dots represent transcripts that are present in the partial annotation. Grey dots represent transcripts that are artificially removed from annotation, i.e., the missing transcripts in the partial annotation. Orange dots represent the unidentified transcript expression for each gene, with each dot representing one gene. Unidentified transcripts are only used for gene expression estimates, but not for transcript expression estimates, leading to improved quantification.

#### 8. Impact of Minimap2 alignment parameters used on NanoCount and Salmon quantification results

For the transcriptome alignment-based methods NanoCount and Salmon, we aligned fastq files to the transcriptome with the recommended/default alignment steps that differ in the number of alignments reported for each read (Leger, 2020; Sipos et al., 2021)(see Supplementary Text Table 3 for the detailed parameter settings). To assess the impact of alignment parameters on quantification, we additionally applied NanoCount with alignments generated using Salmon recommended alignment parameters, and Salmon with alignments generated using NanoCount recommended parameters (Supplementary Text Figure 4). We find that NanoCount quantification results for the spike-in transcripts are similar when different alignment parameters are used (Supplementary Text Figure 4). In contrast, the performance of Salmon is affected, with the recommended parameters providing more accurate results (Supplementary Text Figure 4). For both methods, the recommended alignment parameters give better results, therefore we have kept the different alignment settings for the transcriptome-alignment based methods.

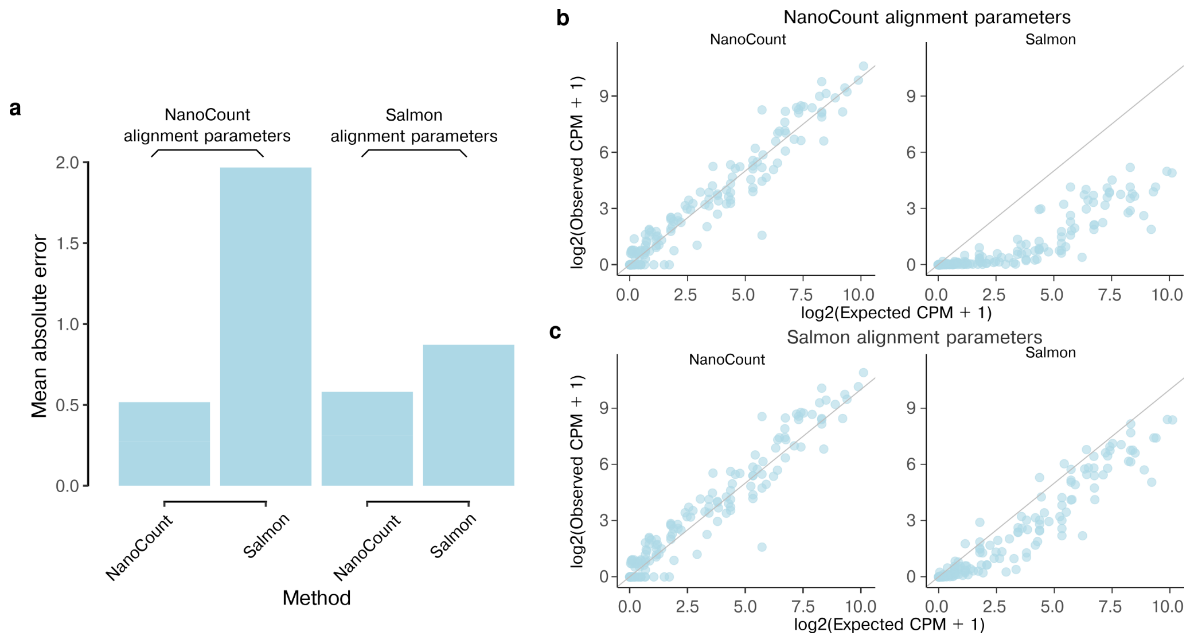


***Supplementary Text Figure 4. The impact of transcriptome alignment parameters on quantification for NanoCount and Salmon***
(a)Shown the barplot of mean absolute error between log2 normalised spike-in transcript abundance estimates and log2 normalised expected abundance when applying NanoCount and Salmon with NanoCount recommended alignments parameters vs with Salmon recommended alignment parameters (b)-(c)Shown the scatterplots of log2 normalised spike-in transcript abundance estimates vs log2 normalised expected abundance when applying NanoCount and Salmon with (b) NanoCount recommended alignment parameters and (c) Salmon recommended alignment parameters are used.

| Method | Minimap2 alignment parameters |
| --- | --- |

| | |
|---|---|
| NanoCount* | " -ax map-ont -p 0 -N 10" |
| Salmon* | "-ax map-ont -p 1.0 -N 100" |
| FLAMES** | "-ax splice -t 12 -k14" and "-ax map-ont -p 0.9 --end-bonus 10 -N 3" |
| All other methods | "-ax splice --junc-bed -k14", "-uf" for stranded samples |

***Supplementary Text Table 3. minimap2 alignment parameters***
*NanoCount and Salmon aligned fastq files to transcriptome fasta file
**FLAMES did alignment twice, once to genome fasta file with "-ax splice -t 12 -k14" and once to transcriptome fasta file with "-ax map-ont -p 0.9 --end-bonus 10 -N 3"

#### 9. Running time and memory usage of Bambu against other methods

Being able to analyse larger sample numbers with reasonable running times was a key consideration during the design and implementation of Bambu. To achieve this, we have vectorised most computation steps for efficient calculation in R, and we have implemented parallel processing and memory friendly file handling. We specifically evaluated the following scenarios for a systematic comparison the running time and memory of Bambu with existing methods for transcript discovery and quantification

(1) Processing of individual samples

For this evaluation, we used 10 samples with varying sequencing depth (500K to 4.5 million reads) that were processed individually using a single CPU. Compared to other transcript discovery methods, Bambu is the second most efficient (average running time of 8.15 minutes and 1.4 GB RAM) after StringTie2 (2.45 minutes, 0.77GB RAM), with all other methods having running times between 14 and 32 minutes (Supplementary Text Figure 5).

For completeness we have also included methods that only perform quantification. As expected, these methods are generally faster as they do not attempt to identify novel transcripts (Supplementary Text Figure 5).

(2) Parallel processing of multiple samples

Most use cases of Bambu will include multiple samples that are jointly analysed. To compare the running time and memory usage of Bambu with other methods in this scenario, we used the same 10 samples and processed them as a single analysis using 5 CPUs. Here, Bambu has a running time of 23 minutes, which is comparable to StringTie2 (21 minutes), and significantly faster than any other transcript discovery method (295-330 minutes, Figure Supplementary Text Figure 5). Bambu supports the re-analysis with pre-processed files (for example when new samples are added to an existing analysis, or alternative thresholds are tested). Using this option further reduces the running time to 14.5 minutes.

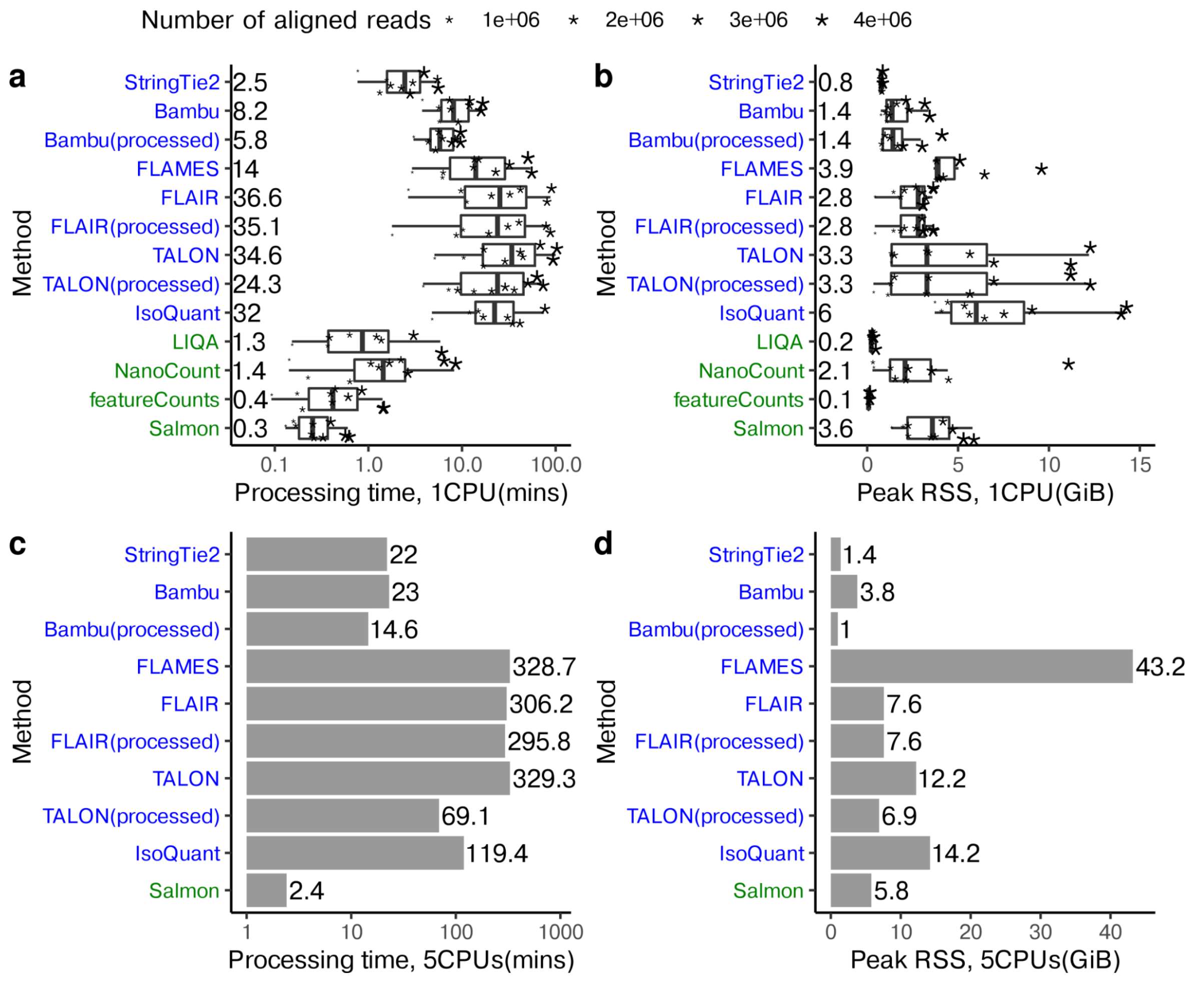


***Supplementary Text Figure 5. Comparison of processing time and peak RSS usage***
Shown is the processing time for 10 samples when (a) processed individually with 1 CPU and (c) processed together with 5 CPUs; and the processing peak resident set size (RSS) usage for 10 samples when (b) processed individually with 1 CPU and (d) processed together with 5 CPUs. Methods that perform both transcript discovery and quantification have names colored in blue, while methods that only perform transcript quantification have names colored in green. Note that FLAMES can process multiple samples when fastq files are provided. IsoQuant is unable to process all samples together as not all samples include the spike in chromosome. All other methods were able to process the data. For IsoQuant, we processed the 10 samples individually with 5 CPUs as an approximation.

(3) Ability to process large data sets (121 million reads)

Due to improvements in the sequencing chemistry, an increasing number of reads is expected from long read RNA-Seq data. We therefore also evaluated the ability to perform transcript discovery and quantification on a sample with very high throughput (121 million reads in total, 94.6 million aligned reads). Bambu can successfully analyse such samples (Supplementary Text Table 4), whereas TALON, IsoQuant, NanoCount and LIQA either return errors or have running times of > 4 days.

| | Method | Processing time (mins) | Peak RAM (GiB) | Number of cpus used* |
|---|---|---|---|---|
| Transcript discovery and quantification methods | Bambu | 98.11 | 43.01 | 1 |
| | StringTie2 | 87.80 | 1.84 | 12 |
| | FLAIR | 701.41 | 7.74 | 12 |
| | FLAMES | 1458.87 | 97.63 | 3 |
| Transcript quantification methods | featureCounts | 31 | 0.07 | 1 |
| | Salmon | 4.98 | 12.42 | 24 |

***Supplementary Text Table 4. Processing time and peak memory usage for a very large sample (121 million reads).***
Only methods that completed the analysis are shown.
*different cpus are used here to quickly process the sample. For featureCounts, no multi-thread is allowed. For Bambu multi-threading is most efficient when multiple samples are provided to facilitate memory-efficient compute, here only a single CPU is used. For flames three CPUs are used to prevent large memory usage

(4) Ease-of-use

Efficiency not only implies low running time and memory, but also ease of use. Bambu was designed to simplify the analysis of long read RNA-Seq data, minimising the number of commands and user-defined thresholds to obtain transcript annotation and quantification results. Among the transcript discovery methods, Bambu, FLAMES, and IsoQuant only require a single command to perform transcript discovery and quantification of 10 samples (Supplementary Text Table 5). In contrast, other methods analyse samples individually before combining them, requiring 10 to 12 commands to obtain the results from such a multi-sample analysis (Supplementary Text Table 5). Compared to FLAMES and IsoQuant, Bambu is significantly faster (see above).

| **Method** | | **Number of commands for 10 samples (ease of use)** | |
|---|---|---|---|
| | | **discovery** | **quantification** |
| *Transcript discovery and quantification* | *Bambu* | 1 command | |
| | *FLAIR* | 10 Correct + 1 Collapse | Quant |

| | | | |
|---|---|---|---|
| | | (11 commands) | 1 command |
| | *TALON* | 1 Initialise DB + 10 talon_label_reads + 1 discovery (12 commands) | Quant 1 command |
| | *StringTie2* | 10 Discovery + 1 merge (11 commands) | Need to re-run StringTie again with “-B -e” parameters (10 commands) |
| | *FLAMES* | 1 command | |
| | *IsoQuant* | 1 command | |
| *Transcript quantification only* | *LIQA* | Not applicable | 10 commands |
| | *NanoCount* | Not applicable | 10 commands |
| | *featureCounts* | Not applicable | 10 commands |
| | *Salmon* | Not applicable | 10 commands |

***Supplementary Text Table 5. Ease of use in transcript discovery and quantification when processing 10 samples***

#### 10. Feature comparison

In this section, we highlight the novel aspects for quantification of long read RNA-Seq in Bambu and present them in a table (Supplementary Text Table 6).

| Method provide long read quantification | Able to quantify gene expression | Able to process multiple samples | No extra steps need to match annotations | Able to track full-length and unique read count estimates | Able to process very large sample (over 100 million reads, using 96 processors and 186.7GiB RAM) |
|---|---|---|---|---|---|
| Bambu | ✔ | ✔ | ✔ | ✔ | ✔ |
| NanoCount | ❌ | ❌ | ✔ | ❌ | ❌ |
| Salmon | ✔* | ❌ | ✔ | ❌ | ✔ |
| featureCounts | ✔* | ❌ | ✔ | ❌ | ✔ |
| StringTie2 | ✔* | ❌ | ✔ | ❌ | ✔ |
| FLAIR | ✔* | ✔ | ❌ | ❌ | ✔ |
| TALON | ✔ | ✔ | ✔ | ❌ | ❌ |
| LIQA | ❌ | ✔ | ✔ | ❌ | ✔** |
| FLAMES | ✔* | ✔ | ✔ | ❌ | ✔ |
| IsoQuant | ✔* | ✔ | ✔ | ❌ | ❌ |

***Supplementary Text Table 6. Feature comparison of methods that provide transcript quantification for long read data***

✔: Possible

❌: Not possible

* not using incompatible reads;

**The whole process takes about 4 days.
